# Synthetic cell-cell adhesion provides benefits of proximity without diffusion-related costs

**DOI:** 10.64898/2026.05.28.728462

**Authors:** Heidi Klumpe, Sarah Loshinsky, Daniel Hart, Kai Tong, Mary J. Dunlop, Ahmad S. Khalil

## Abstract

Multicellularity provides many advantages over unicellularity, but accessing those benefits with synthetic biology approaches is challenging due to potential tradeoffs. The same diffusion limitations that exclude toxins can limit nutrient transport, and small intercellular spaces that facilitate sharing also increase local competition. To isolate when and how cell aggregation, an essential feature of multicellularity, is useful, we compared yeast strains differing only in expression of a single adhesion gene. Unexpectedly, aggregates containing thousands of cells showed no signs of diffusion limitations, growing as well as unicellular strains in both rich and toxic media, likely because reversible cell adhesion allows cells to exchange positions. Aggregating cells also grew better than unicellular strains in the limiting carbon conditions expected to increase diffusion limitations. In mixtures of different genotypes, these benefits were specific to cells expressing the adhesion protein. Together, these results suggest key design principles for engineering aggregates: reformable bonds provide a mechanism to sidestep the fitness costs of increased size, and the benefits within such groups may arise more from nutrient sharing than from diffusion limitations.

## INTRODUCTION

Aggregation into multicellular groups can provide many advantages over unicellular life.^1^ Cells inside an aggregate are protected from toxic substances in the bulk media and unicellular predators. Cells are also closer together, which facilitates rapid communication and metabolite sharing. This coordination can also support division of labor, where the group benefits from the simultaneous and specialized labor of different cell types. In synthetic biology, increasing the survival, cooperation, and functional complexity of engineered strains is critical, and engineering multicellularity into well-known unicellular chassis, including bacteria, yeast, and mammalian cell lines, represents an emerging strategy to advance the field. Recent work already shows the promise of division of labor between specialized strains^2^ and metabolic cooperation,^3–5^ as well as faster settling to improve cell recovery from bioreactors^6^ and ability to bind pathogenic targets.^7^

However, aggregation is not necessarily as beneficial as its frequent evolution^8–11^ might suggest. A major challenge is poor mass transport, as outer cells consume nutrients faster than they reach cells at the center. Delivery of oxygen,^12–15^ sugars,^16^ and other nutrients^3^ is limited in aggregates as diverse as biofilms and organoids. The associated fitness costs can decrease growth rates, erase the benefits of aggregation, and even lead to a reversion to unicellularity.^17,18^

Engineering synthetic multicellularity therefore requires approaches to understand and then balance fitness tradeoffs (Figure 1A). Attachment mechanisms range from reformable bonds (e.g., cell-cell adhesion proteins or sticky extracellular matrix) to more permanent attachment (e.g., incomplete cytokinesis or shared cell walls),^19^ and different multicellular forms can have distinct fitness consequences. Studying these effects in existing multicellular organisms is challenging, as cell-cell attachment mechanisms are polygenic, redundant, and therefore difficult to genetically modify. The effects of modifying aggregation can also be pleiotropic, as diverse mechanisms depend on cell-cell attachment. Moreover, while early efforts to engineer adhesins have focused on key properties like orthogonality and patterning, they have not explored critical parameters for fitness and mass transport limitations, like aggregate size and nutrient concentration.

**Figure 1.**
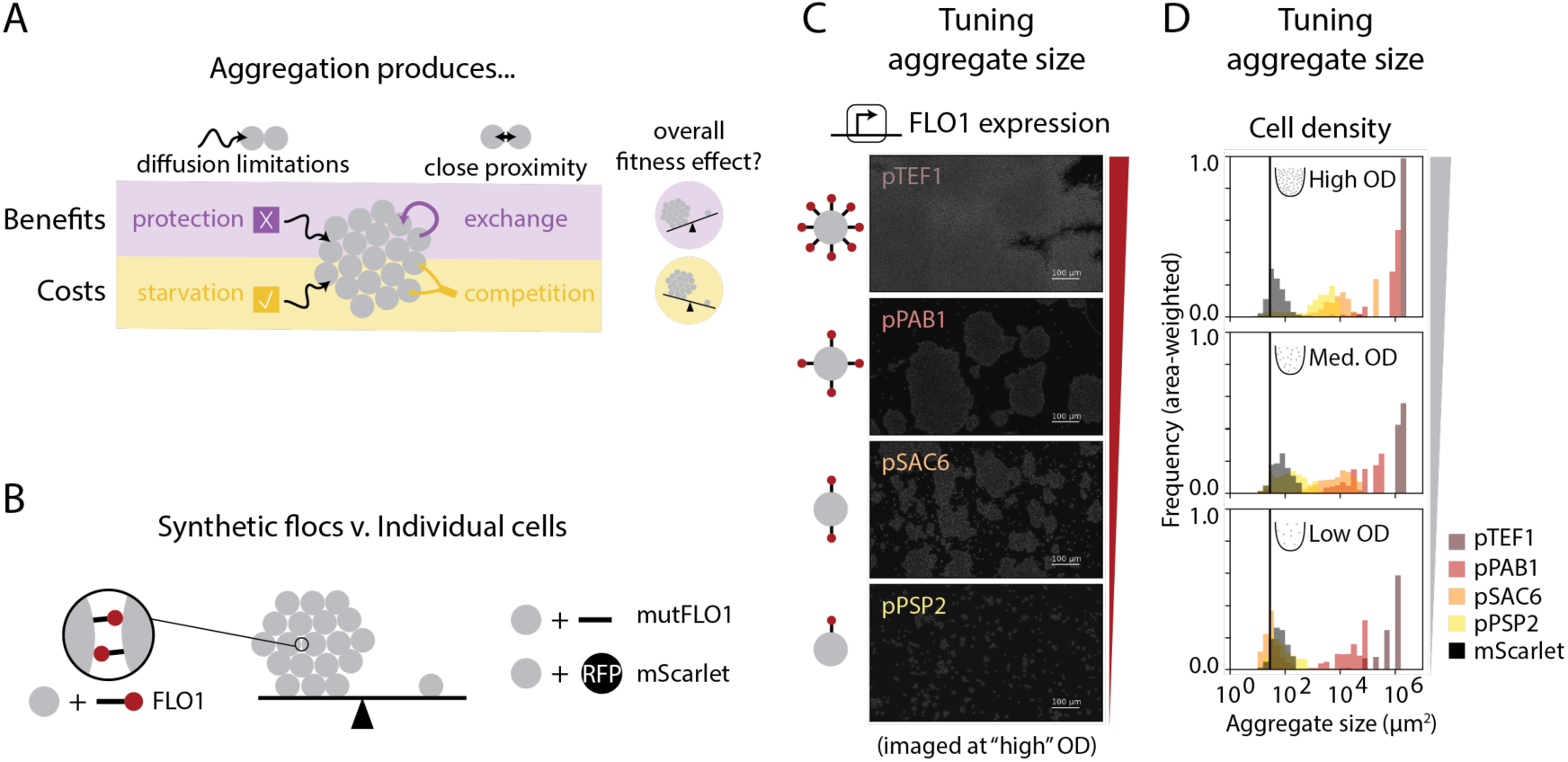
Adhesion proteins provide a model of cell aggregation to study its effects on fitness and function. **(A)** Cell aggregation introduces diffusion limitations and closer proximity between cells. These changes can be both beneficial and costly with respect to the fitness of individual cells. Diffusion limitations of toxic molecules (“X” in the schematic) provide protection, while limited diffusion of nutrients (“✓” in the schematic) reduces growth rates. Close proximity of cells facilitates exchange of nutrients between cells, “but also increases demand per unit volume of media. The net effects of these changes depends on many physical and metabolic parameters, such that the overall fitness benefits of aggregation remain unclear. **(B)** Expressing a single homotypic adhesion protein is sufficient to induce aggregation. As corresponding unicellular controls, either a mutated adhesion protein (e.g., mutFLO1) that could not promote aggregation or a fluorescent protein were expressed in the same non-aggregating base strain. **(C)** Higher FLO1 expression from stronger promoters produces larger aggregates, and the sizes of aggregates range across multiple orders of magnitude. Microscopy images are aggregates from overnight cultures flattened between a coverslip and glass slide into a 2-D monolayer. Distributions are clipped at a maximum size set by the size of the image, which is larger than what is shown in panel B. **(D)** Higher density culture, where collisions between cells are more likely, also produces larger aggregates.

To address this gap, we developed a synthetic system to study the effects of cell aggregation on cell fitness and function. We focused on the reversible bonds of adhesion proteins, as they are a promising tool for engineering multicellularity as well as highly prevalent across prokaryotes^20^ and eukaryotes.^21^ Yeast flocculation, which uses homotypic cell-cell adhesion, makes it possible to study aggregation in a single strain and genotype, a key advantage over other heterotypic synthetic systems. Prior work has demonstrated that inducing expression of flocculation proteins produces macroscopic aggregates from an otherwise unicellular strain, making it possible to study how fitness varies across different length scales. Additionally, prior work suggested that flocculation represents a tradeoff between the costs of expression and the benefits it provided in the presence of ethanol,^22,23^ a condition that can be quantitatively controlled in laboratory evolution experiments. Moreover, *S. cerevisiae* are commonly used in such evolution experiments, including to model the evolution of multicellularity.^24^ Lastly, understanding the effects of aggregation in yeast is relevant to its applications in bioremediation and biomanufacturing, where aggregation can improve stress tolerance and metabolite sharing.

We therefore constructed synthetic flocs by engineering expression of flocculation genes in *S. cerevisiae*. Though we anticipated diffusion limitations to negatively impact fitness, macroscopic synthetic aggregates grew at the same rate as unicellular controls and stably aggregated over hundreds of generations. Not only were large growth defects absent, but no growth benefits were detected in ethanol, the condition where most wild strains flocculate. Flocculation also failed to protect cells from larger, more slowly diffusing toxins, including a cell-wall degrading enzyme. Though unexpected, the absence of growth defects and the absence of protection were consistent with absent diffusion limitations, where most nutrients and toxins readily entered aggregates. We hypothesize that this is because reversible FLO1 binding allows cells to recycle to the aggregate surface, a key difference from smaller yeast aggregates with nearly permanent bonds where diffusion limitations have been reported.^14^ Synthetic flocs also showed evidence of one strong benefit, as they grew faster than unicellular controls in limiting amounts of diverse carbon sources, potentially consistent with a shared extracellular metabolite. Surprisingly, though flocculation permits non-specific binding, FLO1-expressing cells outcompeted non-aggregating cells in limiting carbon. Overall, FLO1 provides a low-cost mechanism for achieving aggregates of diverse sizes, but its advantages may be specific to nutrient-limited contexts and, despite its permissive binding, privatized to its own genotype. Other approaches allowing reversible binding may have similarly limited effects on fitness and provide specifically metabolic, but not protective, benefits to cells.

## RESULTS

### Flocculation proteins produce aggregates of diverse sizes

To isolate the effect of cell aggregation on fitness and function, we wanted a programmable and genetically tunable model of cell aggregation. Therefore, we constructed our synthetic aggregates from the bottom up by first building a non-aggregating, constitutively unicellular strain and then displaying a single adhesion protein on the cell surface (Figure 1B). For the non-aggregating base strain, we deleted genes that could promote cell-cell adhesion (i.e., the sexual agglutination genes SAG1, AGA1, AGA2, and FIG2; and the endogenous flocculation genes FLO1, FLO5, FLO9, FLO10, and FLO11) from BY4742, a commonly used, haploid *S. cerevisiae* strain.

We induced synthetic aggregation by engineering high expression of different FLO proteins into the non-aggregating base strain (Figure S1A). We chose the FLO protein family as our set of candidates as they are associated with protection from chemical stress,^25^ as well as being homotypic (i.e., requiring only a single gene for aggregates to form), likely to express well in their native host, and easily reversed by chelating calcium ions with EDTA. Moreover, ectopically expressing each FLO gene individually allowed us to understand the effect of each FLO protein in isolation and at sufficiently high expression levels. For parallel unicellular controls, we expressed either a truncated FLO protein that did not produce aggregates (i.e., “mutFLO1” whic lacks the PA14 domain) or only a fluorescent protein (e.g., mScarlet).

Despite the similarity of their structures, not all FLO proteins produced aggregates (Figure S1A). While high expression of FLO1 and FLO5 produced macroscopic aggregates visible to the naked eye, high expression of FLO10 and FLO11 did not produce any aggregates, which we confirmed by imaging aggregates flattened between a glass slide and cover slip (Figure S1A). The absence of any effect for FLO10 expression was surprising, given that FLO1, FLO5, and FLO10 are equivalent lengths and rely on a similarly-annotated PA14 domain. However, FLO10’s PA14 sequence is only 75% identical or chemically similar to that of FLO1, and small allelic differences, outside the adhesion domain and especially for FLO11, have been reported to affect overall flocculation.^25–27^

Heterotypic synthetic adhesins can also be co-expressed to produce a similar, single genotype aggregate, but co-expression of heterotypics pairs in a single strain reduced aggregate size considerably, when compared to co-cultures of strains expressing one of the two heterotypic parts (Figure S1B). Changing the display domain also influenced aggregate size, which, similar to the effects of changing FLO1’s stalk,^25^ underscores the importance of sequences outside the adhesin domain for determining overall aggregation. We ultimately chose to continue with FLO1 for our synthetic flocs, as it was the simplest design and produced the largest aggregates.

Controlling aggregate size is a valuable tool for exploring how costs and benefits of aggregation vary across different length scales. We hypothesized that decreasing FLO1 expression would reduce aggregate size by changing overall avidity of cells for each other. We therefore tuned FLO1 expression using either different strength constitutive promoters from the yeast toolkit (YTK)^28^ (Figure 1C) or small-molecule inducible control^29^ (Figure S1C). Both approaches achieved similar levels of FLO1 expression, as reported by co-transcribed mScarlet (Figure S1D), and were quantified at similar culture densities (Figure S1E). Higher FLO1 expression indeed produced larger aggregates, and aggregate sizes could be tuned across multiple orders of magnitude, ranging from tens to thousands of cells per aggregate. Here, we used the same imaging technique as before, carefully and quickly transferring aggregates to a glass slide before flattening with a coverslip, as the non-covalent attachments between cells were perturbed by vigorous pipetting or long-term settling on the bench. As expected, the unicellular controls did not aggregate (Figure S1C).

We further hypothesized that denser cultures have higher probabilities of cell collisions and therefore maintain larger aggregate sizes. Indeed, cultures initiated at higher OD had larger aggregates (Figure 1D, S1C), whose area we quantified by thresholding on a constitutive fluorescent reporter. In this way, similar size aggregates could be achieved by different FLO1 expression levels, but at different culture densities. Thus, gene expression level and cell density are both crucial parameters for tuning aggregate size.

### FLO1-based aggregation has little effect on growth rate

Higher consumption of nutrients and their decreased diffusion into an aggregate can reduce the growth rate and overall fitness of aggregates^16^ (Figure 2A). Such diffusion limitations are frequently observed in biofilms, tumors, and organoids, in particular for oxygen.^12,13,15^ Prior studies of flocculating strains have also reported slower glucose consumption^23,25^ and reduced growth rates. Specifically, a strain with galactose-induced FLO1 expression grew four-fold slower than a FLO1 knockout in the same conditions, with similar differences in growth between flocculating and non-flocculating variants of a wild strain.^22^

**Figure 2.**
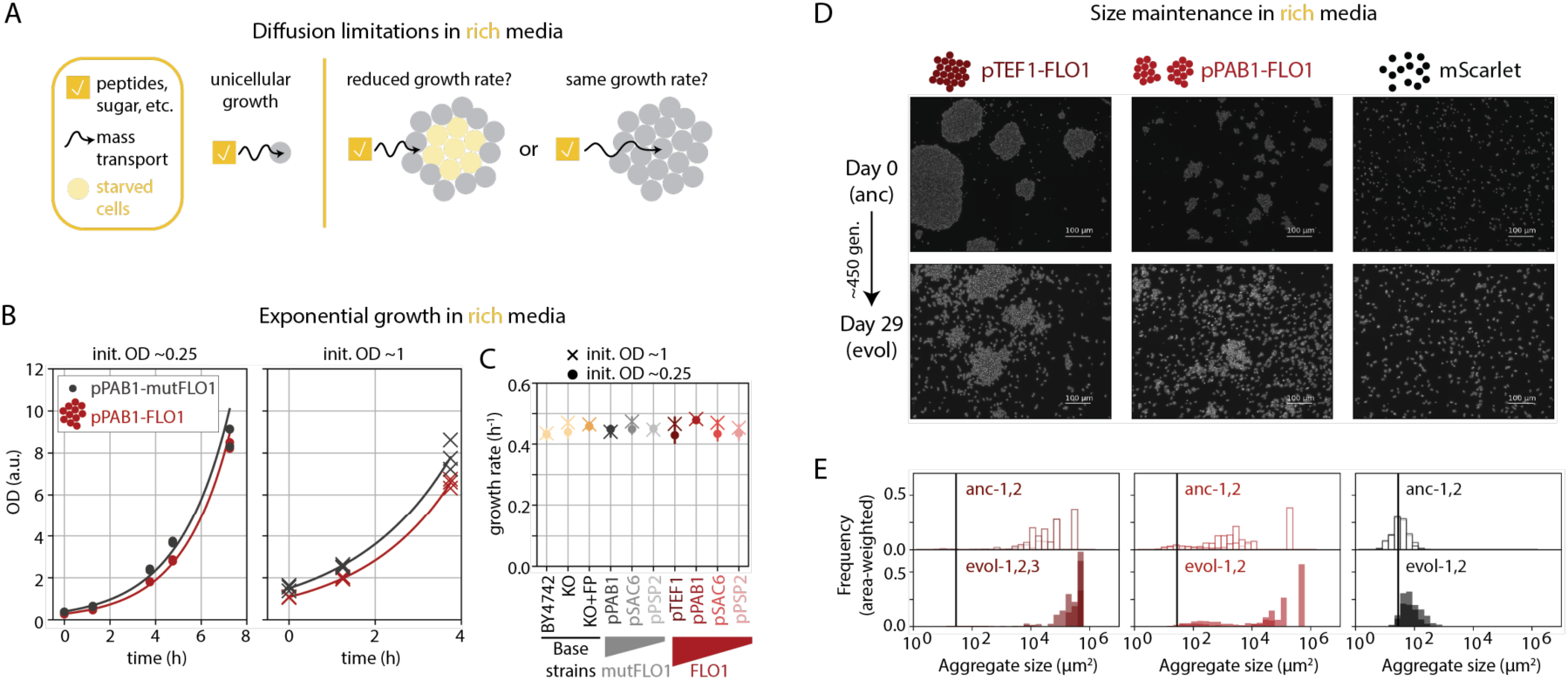
FLO1 expression has no detectable effect on growth rate and is stably expressed over hundreds of generations. **(A)** One possible fitness cost of aggregation is the reduced mass transfer of key nutrients, especially when grown in rich media like YPD. This would manifest as a reduced growth rate of aggregates relative to unicellular controls. **(B)** Strains expressing either FLO1 or mutFLO1 from the same promoter have similar exponential growth rates in YPD. This is true across the aggregate sizes achieved at lower (0.25) and higher (1.0) initial ODs. Datapoints are independent wells grown from the same dilution of overnight culture and sacrificially measured at the timepoint indicated. Lines are the least squares fit of exponential growth. **(C)** Growth rates of diverse aggregating and non-aggregating strains were measured by fitting exponential growth rate curves to similar OD dynamic measurements as in panel B. These include various base strains with different auxotrophies, including the parental BY4742 strain, the FLO and sexual agglutination gene knockout, and the knockout strain expressing two fluorescent proteins (mNeonGreen and mTagBFP2), as well as strains expressing mutFLO1 or FLO1 from different strength promoters. Different marker styles indicate different initial ODs, as in panel B. Error bars are the square root of the variance for the growth rate estimate. **(D)** Aggregating strains expressing FLO1 from a strong (pTEF1) and medium (pPAB1) strength promoters were continuously cultured at exponential phase using the eVOLVER platform, alongside a control unicellular strain with mScarlet under control of pTEF1. After 29 days of continuous growth in YPD media, samples from each vial (“evol”) were imaged as a monolayer between a glass slide and cover slip, alongside ancestral parental strains (“anc”) grown overnight in YPD. **(E)** Aggregate sizes in multiple fields of view of both the evolved (“evol”) and ancestral (“anc”) strains were quantified by thresholding on constitutively expressed mNeonGreen in all strains. Replicates of the ancestral strain (e.g., anc-1, anc-2) were different overnight cultures of the same strain, while replicates of the evolved strain (e.g., evol-1, evol-2) were different vials on the eVOLVER, with an additional vial for the largest aggregates. The frequency of each aggregate size is weighted by the aggregate size, to estimate the probability that a single cell from the population appears in an aggregate of a particular size. Aggregate size is largely maintained across the many weeks of the experiment, suggesting aggregation does not significantly affect fitness.

To quantify any such large negative effect on fitness in our strains, we measured growth across different unicellular controls and synthetic flocs in YPD, the standard rich media for yeast culture (Figure 2B). We sampled cultures approximately every two hours, broke up flocs by adding EDTA, and measured OD. As unicellular controls, we included the mutFLO1 and mScarlet unicellular strains, as well as the wildtype background (BY4742), the non-aggregating base strain (KO), and a derived fluorescent strain (KO+FP). Because diffusion limitations may occur at a particular (i.e., sufficiently large) size, we varied aggregate size by changing two variables: FLO1 expression (cf. Figure 1C) and initial culture density (cf. Figure 1D). Based on quantification of the largest synthetic flocs (i.e., pTEF1-FLO1, cf. Figure 1D and S1F), we estimate the median aggregate radius begins around 50μm at an OD of 2 and increases to at least 100μm at an OD of 6 (see supplementary note).

However, growth rates did not vary considerably between strains or initial ODs, as the estimated ranges for best-fit parameters were overlapping (Figure 2C). Similar growth among background strains suggested the deletion of native FLO genes and the ectopic expression of fluorescent proteins did not have dramatic effects on growth rates. The limited effects of expressing mutFLO1 suggests that the cost of expressing FLO1, a protein of similar size and composition, is small, consistent with a 3% fitness difference associated with FLO1 expression determined from a competition assay.^22^ Most notably, synthetic flocs had the same growth rates as all other strains tested, even for larger aggregates formed at higher OD.

The absence of a large growth defect in synthetic flocs was unexpected, particularly given the prior report of four-fold reduced growth in a similar BY4741 background, the same FLO1 coding sequence, and rich YPD media. However, we hypothesize that the key difference between these experiments is aggregate size, as the reduced growth rate appeared in millimeter-scale flocs much larger than ours. Such drastically larger sizes may be necessary to produce diffusion limitations. Another study of flocs in bioreactors proposed 100µm as the lower bound at which diffusion limitations first appear,^30^ though in those experiments, larger flocs were produced by slower stir speeds, and reduced convection could also explain some of the slower glucose transport.

Nonetheless, in our experiments, FLO1 expression has no obvious effect on growth rate across a range of culture densities and expression levels relevant for many synthetic biology experiments. While glucose diffusion limitations may still occur at larger aggregate sizes, this may be an extreme parameter regime. For example, constructing a 2mm synthetic floc would require approximately every cell in 3mL culture already at an OD of 6 (see supplementary note). In addition, two studies of flocs in bioreactors that reported slower glucose consumption only inoculated flocs at high cell densities, explicitly using long preculturing and larger volumes to increase aggregate sizes, as well as supplementing additional calcium ions that could further promote flocculation.^25,30^ Not only are such sizes potentially hard to achieve, but they require nearly saturated cultures, where the rapid consumption of nutrients slows growth of most strains independent of flocculation. Overall, our results indicate that for a wide range of culture conditions and FLO1 expression levels, synthetic flocs can have similar growth rates as unicellular strains.

### Synthetic flocs are evolutionarily robust

While our growth measurements exclude any large fitness defects, smaller and harder-to-detect fitness costs could increase the chances of reverting to unicellularity, as non-aggregating mutants that do not experience the same costs can outcompete the aggregating cells.^14,17,18,31^ For example, in *S. cerevisiae*, aggregates held together by incomplete cell separation (another aggregation mechanism) have reverted to smaller sizes in continuous culture,^18,32^ and loss of flocculation during long-term growth in rich media has been observed.^33^ Such vulnerability to reversion would pose a significant challenge to engineered multicellularity.

To test the long-term robustness of FLO1 expression and flocculation, we continuously cultured synthetic flocs in rich media using the eVOLVER platform, a device for automated continuous culture.^34^ We used the parallel vials on the eVOLVER to culture multiple biological replicates of three strains: high FLO1 expression (pTEF1), medium FLO1 expression (pPAB1), and the mScarlet unicellular control. We set the eVOLVER to maintain each culture in a narrow OD window around 0.7. While the OD of aggregating strains is only approximate, maintaining cultures in exponential phase maximized the number of generations reached during the experiment, and the frequent small dilutions (e.g., replacing only a fraction of the 20mL culture volume) kept aggregate size relatively constant over time. To monitor aggregate size and FLO1 expression, we imaged flattened aggregates from cultures backdiluted to the same OD and measured levels of co-transcribed mScarlet with flow cytometry.

At the end of four weeks, or approximately 450 generations, the “evolved” strains had the same distribution of aggregate sizes as the initial cultures (Figure 2E). Aggregates appeared to get slightly larger over the course of the four weeks, but this was due to an increase in cell size and aspect ratio that occurred in all vials (Figure S2A,B). Cells also continued to express mScarlet, suggesting FLO1 expression was also maintained (Figure S2C). In fact, all vials, including the non-aggregating controls, showed a slight increase in mScarlet. This increase of both mScarlet and cell size suggested an increase in ploidy, which is commonly observed in experimental evolution of haploid yeast^35^ and which we confirmed by propidium iodide staining (Figure S2D,E). Thus, even in the midst of other genetic adaptations to growth on rich media, flocculation was stably maintained for hundreds of generations of growth, consistent with the similar growth rates measured in rich media (Figure 2C).

Maintained FLO1 expression was somewhat unexpected given a prior observation of flocculation being lost,^33^ but we again hypothesize that this is due to differences in the culture density and overall aggregate size, as the background strain, FLO1 gene, media composition, and number of generations were otherwise similar. The other long-term culturing experiment used daily 100-fold dilutions, in contrast to the eVOLVER’s small dilutions every two hours. As flocs can double at least 1000-fold in a day, we predict these cultures were saturated for much of the experiment. The resulting larger aggregate sizes and reduced glucose in the media could both increase the penalty for flocculation, increasing the probability of reversion. We therefore hypothesize that either limiting aggregate size or time at saturation could be sufficient to ensure long-term FLO1 expression.

### High culture density, but not FLO1 expression, protects cells from ethanol and other toxins

Besides excluding nutrients, aggregation can protect cells by shielding the aggregate center from toxic molecules (Figure 3A), as observed in the increased antibiotic resistance of biofilms.^36,37^ Many wild yeasts flocculate as a response to ethanol, and flocculation has been associated with protection from organic acids (e.g., acetic and lactic acid),^38,39^ ethanol,^22,40^ bioethanol inhibitors (e.g., furfural),^25,41^ and fungicide.^22^ Because such protection can come directly from diffusion limitations, we wanted to achieve the largest aggregate sizes possible. We therefore used the synthetic flocs with inducible FLO1 (cf. Figure S1B) which could achieve the highest FLO1 expression (Figure S1C).

**Figure 3.**
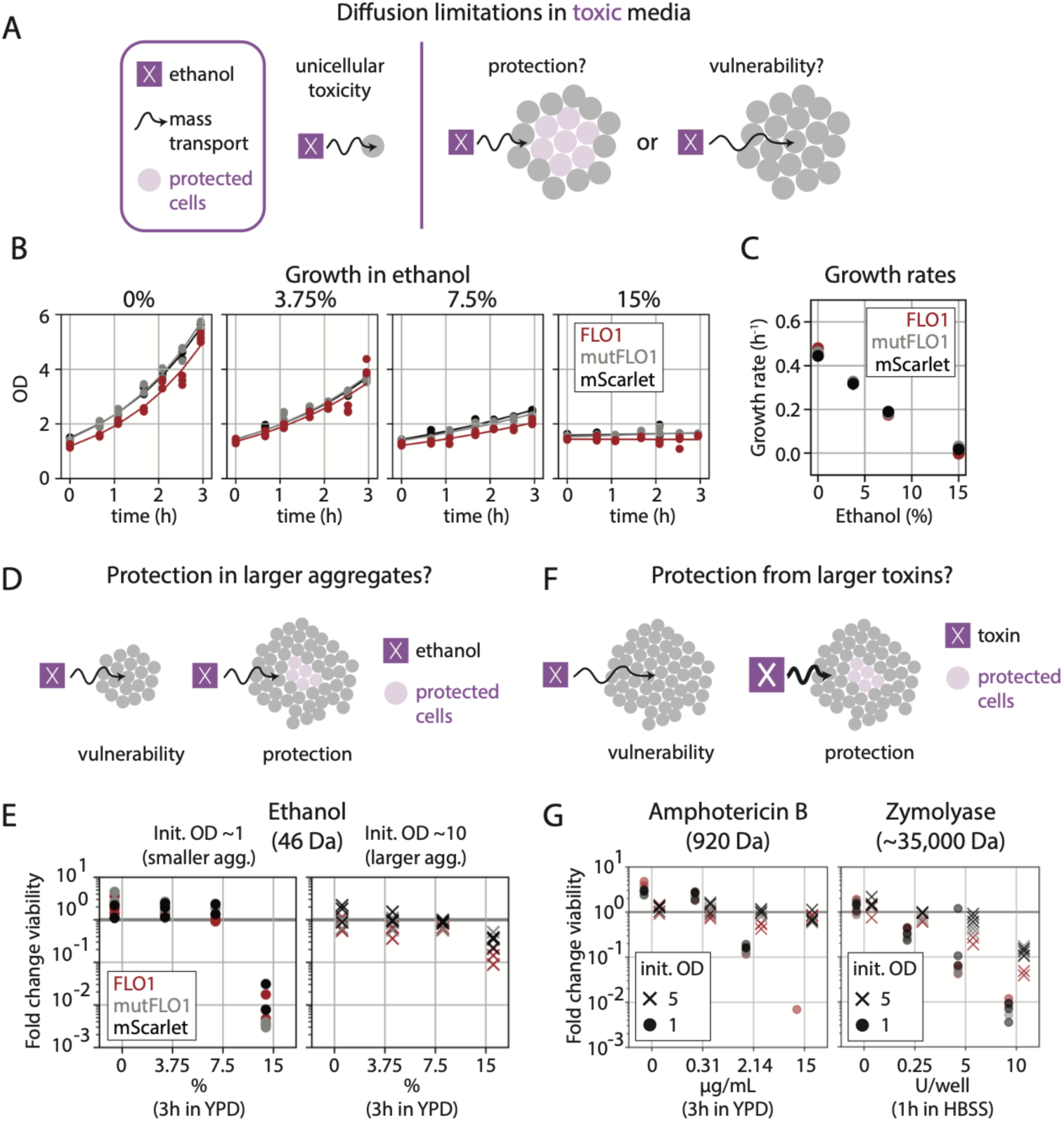
Ectopic FLO1 expression does not protect cells from diverse chemical stresses. **(A)** High concentrations of ethanol are toxic to individual cells, but diffusion limitations in an aggregate could protect an inner core of cells. **(B)** Triplicate cultures of synthetic flocs, the mutFLO1 strain, and the mScarlet strain (all inducible and maximally induced) have similar growth rates across many ethanol concentrations. Lines indicate the best fit exponential growth curve. **(C)** Growth rates decrease as the ethanol concentration increases, but equally for all strains and regardless of FLO1 expression. Error bars are the square root of the variance of the growth rate estimate. **(D)** Larger aggregates may provide better protection from ethanol because diffusion limitations can arise at larger length scales. **(E)** Triplicate cultures of the same strains as in panel B were initiated at high and low OD before adding ethanol, and viability was estimated by plating countable colonies. At low initial OD, cells continue to grow and divide in up to 7.5% ethanol, but die rapidly in 15% ethanol. At higher initial OD, cells do not grow because they are near saturation and are less affected by 15% ethanol. This protective effective is similar for all strains. **(F)** Aggregates may provide protection from larger toxins, as a larger molecular weight can decrease diffusivity and therefore produce diffusion limitations. **(G)** Amphotericin B (∼20x the molecular weight of ethanol) and Zymolyase (∼750x the molecular weight of ethanol) decrease the viability of both unicellular and aggregating strains. Similar to ethanol, toxicity increases with toxin concentration, but higher initial OD provides a protective effect.

We first measured growth in ethanol, which induces flocculation in wild strains. We exposed cells to a range of ethanol concentrations, reasoning that higher concentrations would have larger effects on growth but lower concentrations were more likely to be diffusion-limited. We included the inducible versions of both unicellular controls, mutFLO1 and mScarlet, and cultured cells in a shaking incubator for two hours before ethanol was added, to ensure aggregates were formed. However, ethanol reduced growth of all strains equally (Figure 3B,C). We also counted colony forming units to check for losses of viability not captured by OD. Indeed, 15% ethanol reduced the number of viable cells (Figure S3A), though the OD remained apparently constant. Nonetheless, this loss of viability was the same for both synthetic flocs and unicellular controls, confirming that FLO1 provided no protection from ethanol.

We hypothesized that, at the start of the experiment, flocs may not have been large enough to protect cells from ethanol. Because we had already maximized FLO1 expression, we generated larger aggregates by starting the experiment at a higher initial culture density, near saturation (Figure 3D). As OD does not change much in these nearly saturated cultures, we focused on counting colony forming units. However, even at higher initial OD, the effects of ethanol were again the same for all strains, with increasing ethanol producing the same decrease in viability (Figure 3E, S3A). Instead, the most notable difference was between high and low culture density. Though 15% ethanol decreased viability in all conditions, higher OD cultures were less affected than cultures at lower OD. Thus, this apparent protective effect was not specific to synthetic flocs but was instead specific to high density culture.

Thus, only culture density, but not FLO1 expression, protected cells from the toxicity of ethanol. This could be because the effective concentration of ethanol per cell is lower (an analogy with the inoculum effect^42^), or because some of ethanol’s toxic effects are growth rate dependent, such as interactions with the replication fork,^43^ and therefore less toxic to slowly growing cells. Given that the conditions of our experiment, with abundant sugar and continuous mixing, are far from the conditions in which flocculation evolved in wild yeast, this result does not necessarily indicate that wildtype flocculation does not protect cells from ethanol. However, it does show that bringing cells together with adhesion proteins, even into very large aggregates, may not be sufficient to provide physical protection from chemical stress, which is relevant for synthetic aggregates.

We next hypothesized that larger toxins, which diffuse more slowly, would be more likely to be excluded from aggregates (Figure 3F). We therefore tested two chemical stressors of larger molecular weights. Amphotericin B is a fungicide of approximately 920 Da (i.e., twenty times more massive than ethanol) and previously reported to be less lethal in flocs than in unicellular cultures.^22^ We subjected the synthetic flocs and unicellular controls to various concentrations of amphotericin B, and the drug affected all strains to the same extent across all doses (Figure 3G, S3B). Higher drug concentrations reduced viability, with almost no colonies observed after three hours of the highest dose. However, viability did not change when cultures started at a near saturating density, even at the highest drug dose. Thus, as with ethanol, higher culture density apparently protected cells, while FLO1 expression had no effect.

For the largest toxin, we used Zymolyase, a proprietary and commercially available enzyme mixture that can kill yeast by degrading the cell wall. We estimate that the molecular weights of the components are similar to related enzymes, such as b-1,3-glucanase from *Klebsiella pneumonia* (5GY3) and a laminase from *Paenibacillus barengoltzii* (5H9Y), which are 35 and 50 kDa respectively. We anticipated that these enzymes would less effectively degrade the cell wall in rich media, either because the yeast extract in YPD provides an alternative substrate for the enzyme or because pH changes in the unbuffered solution affect enzyme activity. We therefore diluted cells in Hanks Balanced Salt Solution (HBSS) before adding the enzyme. Zymolyase reduced cell viability in as few as 30 minutes (Figure S3C) and had similar effect as the other toxins. Higher concentrations decreased viability more, but only higher culture density improved survival (Figure 3G).

Together, these results show that even though synthetic flocs approach a diameter of 100µm, this increased length scale is not sufficient to provide physical protection, even for larger and presumably slower-diffusing toxins. Only increasing culture density provided some protection, but this was independent of flocculation.

### Culture density, but not FLO1, affects gene expression

Despite the absence of strong phenotypes like growth rate defects or chemical protection, flocculation may still affect cell state in subtle ways, such as cell wall remodeling^40,41^ or changing expression of specialized genes.^25,44^ Therefore, performing RNA-Seq to quantify gene expression could both reveal what, if any, biological processes were affected (Figure 4A).

**Figure 4.**
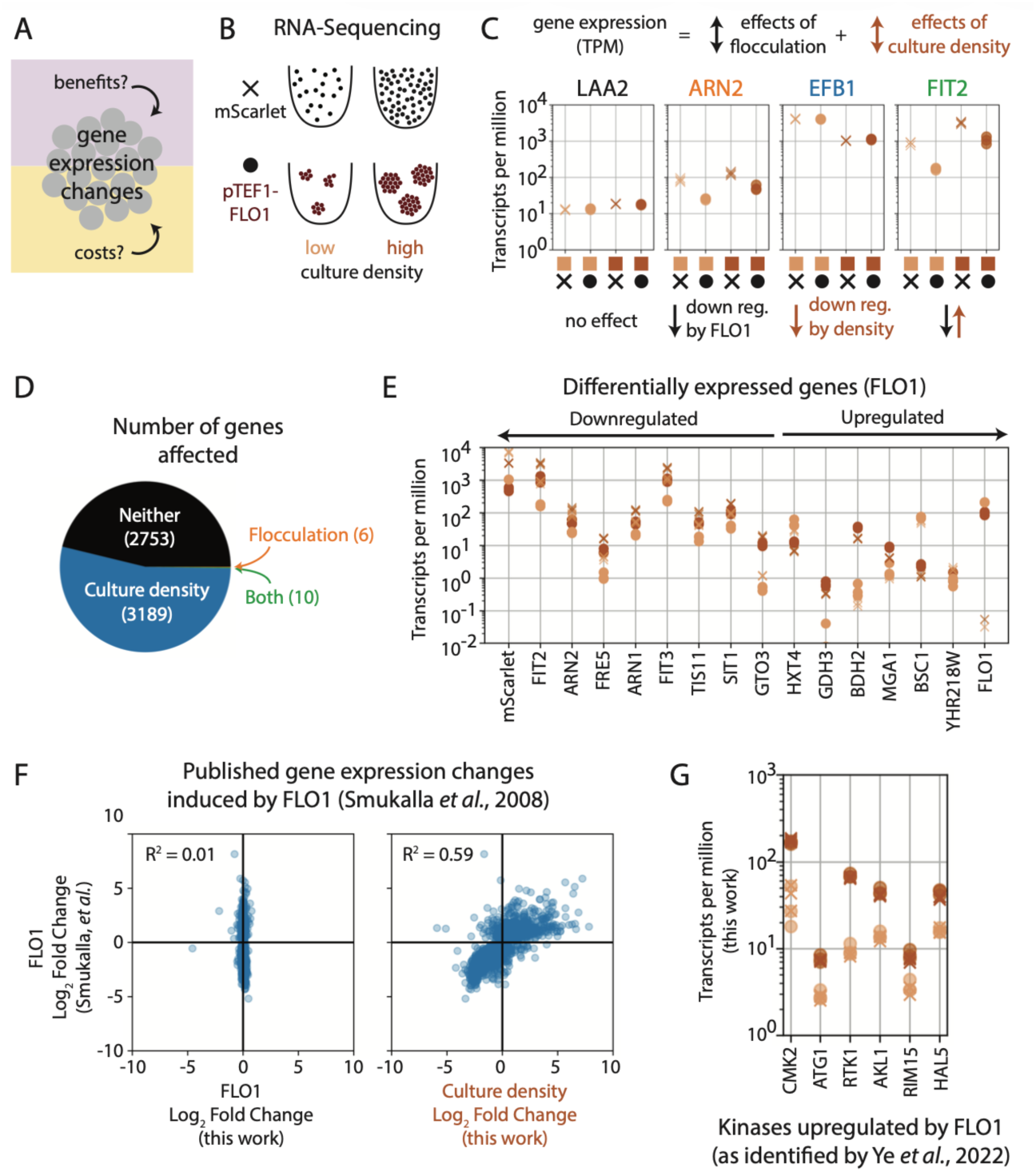
Culture density, but not FLO1 expression, reproduces previously reported effects of flocculation. **(A)** Though large effects on growth rate were absent in extreme culture conditions, aggregation could impact gene expression. **(B)** To generate samples for RNA sequencing, cultures of the unicellular strain (mScarlet) and the larger synthetic flocs (pTEF1-FLO1) used in the long-term culturing experiment (cf. Figure 2) were grown at the same “low” density of exponential phase (∼1) and high density of saturated culture (∼7) in YPD. **(C)** For each gene, comparing its expression across the four samples reveals the effects of flocculation and density. Expression can either be the same in all conditions (e.g., LAA2), change with FLO1 expression but not density (e.g., ARN2), change with density but not FLO1 expression (e.g., EFB1), or change with both density and FLO1 expression (e.g., FIT2). **(D)** After thresholding for effect size and statistical significance, about half of all genes in the transcriptome were identified as sensitive to culture density, while a minute fraction were sensitive to FLO1 expression. **(E)** The sixteen genes identified as sensitive to FLO1 expression included the two integrated genes (mScarletI and FLO1), which had different expression levels in the two strains by design. The genes are sorted by FLO1’s effect size, showing that the most upregulated and downregulated genes are actually FLO1 and mScarlet, respectively. The remainder, while statistically significant, had very small effect sizes. **(F)** Previously reported FLO1-responsive genes^22^ were not sensitive to FLO1 in our dataset. Instead, the reported FLO1 effect sizes correlated with the effect size of culture density. **(G)** Prior work identified genes upregulated in wild flocculating strains^38^ that were responsible for protection from acetic acid. In our samples, these genes are upregulated by culture density and unaffected by FLO1.

We collected RNA from the largest synthetic flocs (pTEF1 promoter, cf. Figure 1C) and the mScarlet unicellular control. We sampled cultures at the same moderate OD of the long-term culture experiment (cf. Figure 2) but also at the nearly saturating OD where aggregates are largest (Figure 4B) by sampling 5h after a 10- or 3-fold (respectively) backdilution. To identify genes whose expression was specifically dependent on flocculation, we modeled gene expression as a linear sum of the effects of flocculation, culture density, and interactions between these two variables (Figure 4C). We then identified genes for which the effects of either flocculation or culture density were both statistically significant (i.e., adjusted p-value < 0.05) and biologically significant (i.e., fold change greater than 1.5) (Figure S4A).

Over half the transcriptome was differentially expressed across the four conditions, but virtually all of these were affected by culture density alone (Figure 4D), which is consistent with widespread changes in gene expression as cultures become saturated.^45^ By contrast, only a fraction of a percent of the transcriptome was specifically affected by flocculation, including the ectopic copies of FLO1, which is more highly expressed in the flocculating strain, and of mScarlet, which has slightly more expression in the reporter than from the co-transcribed FLO1 construct (Figure 4E). In fact, these two genes comprised the largest effect sizes for upregulation and downregulation, respectively, while the remainder of the effects of FLO1, though statistically significant, rarely produced fold-changes larger than 2. However, the downregulated genes were enriched in GO terms related to iron transport and homeostasis (Figure S4B). Many of these genes encode proteins that, like FLO1, are displayed on the cell surface, suggesting possible interactions at the cell wall.

This result is in marked contrast with a previous study that identified over 2000 differentially expressed genes between galactose-induced flocs and a FLO1 knockout in BY4741.^22^ Of these, only two, BDH2 and GDH3, were also identified as FLO1-responsive in our analysis. By contrast, most (i.e., 1697 of the 2165) of the differentially expressed genes in that report were instead significantly affected by culture density in our dataset. In fact, the reported size and direction of FLO1’s effect correlated with the size and direction of culture density’s effect in our dataset (Figure 4F). Another study showed that, when exposed to acetic acid, a wild flocculating strain had higher expression of stress kinases than the same strain with FLO1 knocked out.^38^ However, in our data, both high density cultures, even in the absence of FLO1, increased the expression of these genes (Figure 4G). Because these stress kinases were also found to confer protection from chemical stress, this could explain the protective effect of culture density we observed (cf. Figures 3E,G).

At a minimum, these results show that our approach to engineering synthetic flocs has limited effects on gene expression, consistent with no large effects on growth rate or survival in ethanol. Curiously, we found that previously reported effects of flocculation instead match the effects of culture density in our dataset. Because high culture densities are necessary to produce the large aggregates used for some studies, this is a potentially confounding effect. The study in BY4741 collected samples for transcriptomic analysis after 24 hours of growth, where saturation could indeed have confounding effects on gene expression.^22^ At the same time, the study with the wild strain explicitly collected cells in the exponential phase, suggesting a genotype-specific effect.^38^ At the same time, culture density and flocculation could have similar effects on gene expression as they are both increases in cell density, though either globally or locally.

### Molecules of diverse sizes readily infiltrate flocs

The similarities in growth between synthetic flocs and unicellular strains suggests that cells in these aggregates have ready access to molecules in the bulk media. To more directly measure this, we added cell surface dyes to shaking cultures of synthetic flocs or unicellular controls and measured what proportion of the cells were stained, as a proxy for which cells were exposed to the dye. As before, we hypothesized that larger dyes were more likely to be diffusion-limited (Figure 5A), so we used dyes of diverse sizes. We stained synthetic flocs with different FLO1 expression levels, as larger flocs should exclude more dye. For more direct comparison with the evolution experiment and RNA-Seq data, we dyed the same three strains at a similar culture density that was used for those experiments (OD ∼1). We exposed cells to a brief one-minute pulse of dye before “quenching” with a ten-fold dilution in YPD, which lowered background staining without significantly lowering on-target dyeing (Figure S5A). Such quenching was necessary as we wanted staining to reflect interactions with liquid media in shaking culture, not incidental staining that occurred when aggregates were broken and centrifuged to prepare for flow cytometry. To quantify the level of background staining that could still occur during sample processing and after dye dilution, we resuspended separate, unstained cultures in the same concentration as the “dilute dye” and measured the degree of staining. We optimized concentrations of all dyes such that “dilute” staining was as low as possible while still having detectable staining in the shaking condition.

**Figure 5.**
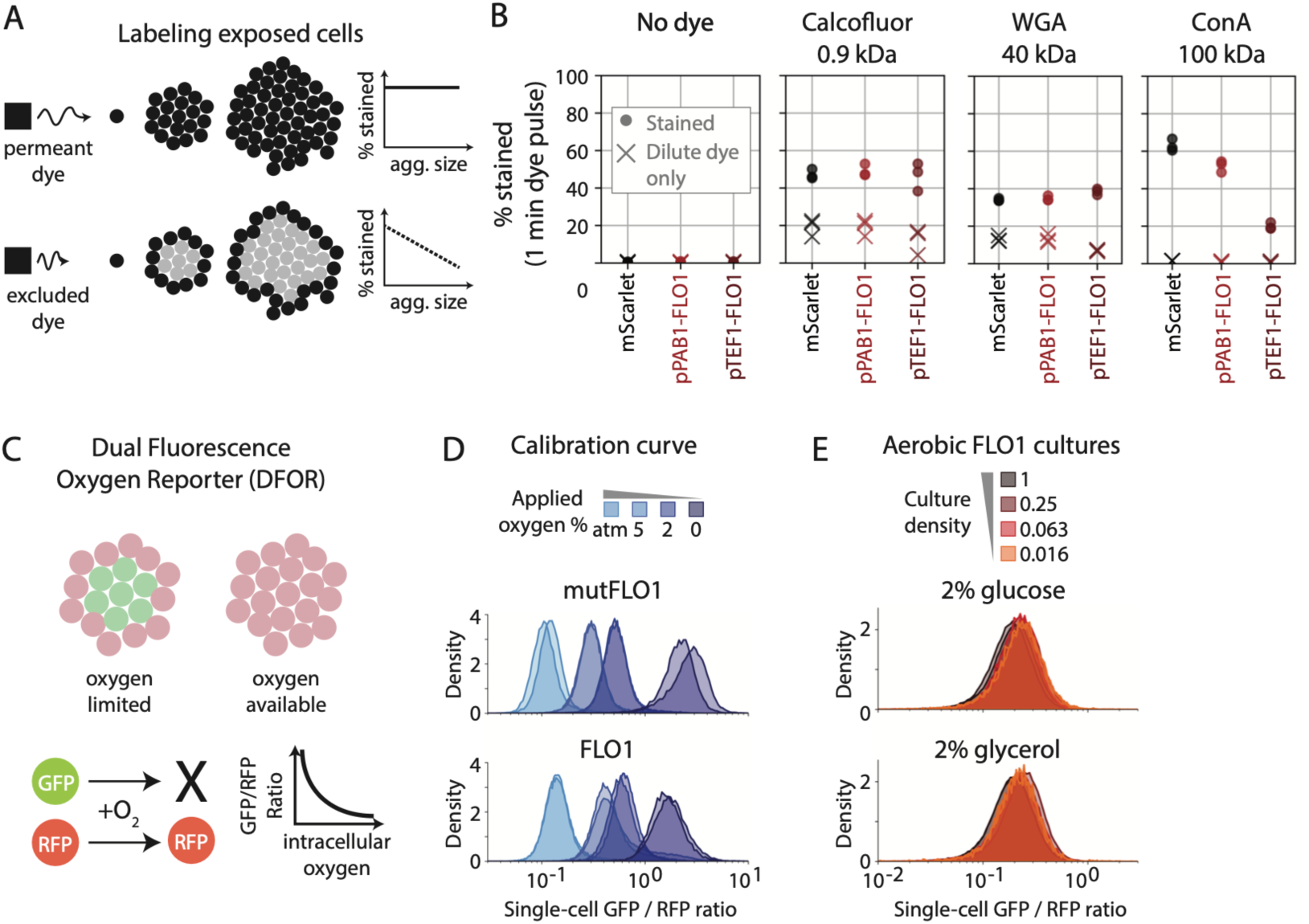
Large dyes and oxygen readily enter FLO1-based aggregates. **(A)** Dyes of different molecular weights can reveal how molecules of diverse sizes enter aggregates, with unequal dyeing of cells if dye is excluded. We hypothesized that while individual cells are exposed equally to all dyes, larger dyes are more likely to be excluded from aggregates. **(B)** The mScarlet unicellular control and synthetic flocs of different sizes were stained with no dye or dyes of increasing molecular weights for 1 minute, before the dye was diluting to stop staining. “Dilute dye” indicates background staining that arises during sample processing, measured with unstained cultures in dilute dye only. Only the largest dye, ConA, was excluded from aggregates. Dyeing was inversely proportional to aggregate size. **(C)** The dual-fluorescent oxygen reporter (DFOR) provides a single-cell readout of intracellular oxygen, another measure of mass transport in aggregates. The ratio of oxygen sensitive GFP to constitutive RFP anti-correlates with intracellular oxygen. **(D)** Using an eVOLVER module with feedback control of oxygen levels, we cultured the synthetic flocs and mutFLO1 control, modified to express the DFOR, at different oxygen levels. We kept culture density low to prohibit large aggregate formation. As expected the ratio of GFP to RFP increased as oxygen was removed. **(E)** Intracellular oxygen concentrations are not reduced as culture density (i.e., aggregate size) increases. The same trend was observed on fermentable (2% glucose) and non-fermentable (2% glycerol) carbon. Thus, flocculation does not appear to limit access to oxygen.

Under this protocol, we stained aggregates with cell surface dyes of three different sizes. Calcoflour, approximately the size of amphotericin B, stained aggregating and non-aggregating cells to the same extent even for larger aggregates (Figure 5B), consistent with equal killing of synthetic flocs and unicellular controls by amphotericin B, as well as cells’ apparently equal access to key nutrients during growth in rich media. Staining with 40 kDa wheat germ agglutinin (WGA) produced the same effect, with all synthetic flocs being stained as well as the mScarlet strain after 1 minute (Figure 5B). This was consistent with the equal effect of Zymolyase whether or not strains expressed FLO1.

Only concanavalin A, a 100 kDa tetramer, had lower staining of synthetic flocs (Figure 5B). Notably, the degree of staining was inversely proportional with aggregate size, consistent with the hypothesis that larger aggregates shield a larger core of cells from the dye. Interestingly, even this exclusion of the largest dye was not permanent. Staining for longer periods of time revealed that ConA dyed all strains equally well after 1 hour (Figure S5B), either due to further diffusion into the aggregates or cell rearrangement due to the reversibility of FLO1 binding. Thus, molecules of diverse sizes can readily enter even the largest synthetic flocs and on time scales relevant for cell growth and death.

### Intracellular oxygen levels are similar across all cells in a floc

While dyeing serves as a proxy for molecules entering an aggregate, it does not fully recreate the consumption by cells that limits mass transfer. Therefore, measuring the concentration of a real metabolite can provide better insight into aggregate permeability. Oxygen is often limiting in cell aggregates, and in snowflake yeast that are held together by nearly permanent bonds, only an outer periphery about three cells thick performs aerobic respiration when cultured on a non-fermentable (i.e., oxygen requiring) carbon source, suggesting oxygen does not penetrate the aggregate when oxygen demand is high.^14^

To quantify intracellular oxygen, we used the fluorescent version of the Dual Luminescence Oxygen Reporter^46^ (i.e., DFOR) for single-cell measurements. In DFOR, degradation of a tagged GFP is proportional to the concentration of oxygen, due to a co-expressed plant enzyme. Because the GFP is co-translated with a stable RFP, the ratio of these fluorophores provides a readout of oxygen concentration (Figure 5C, Figure S5C). We integrated the DFOR into our largest constitutive synthetic floc (i.e., pTEF1-FLO1) and its corresponding mutFLO1 unicellular control, but removed the co-transcribed mScarlet used elsewhere as it overlapped with the DFOR readout.

To verify that the reporter was indeed sensitive to intracellular oxygen, we leveraged a recently developed eVOLVER module that allows control of oxygen levels across 16 vials. We cultured the synthetic flocs and mutFLO1 unicellular controls in different concentrations of oxygen and used turbidostat control to maintain cultures at low density (OD ∼0.3), which we visually confirmed limited aggregation. After blocking translation with cycloheximide, we removed cells from the eVOLVER and measured single-cell levels of RFP and GFP by flow cytometry.

As expected, the ratio of GFP to RFP increased as oxygen decreased (Figure 5D). For both strains, the distribution of single-cell values was unimodal, reflecting both the homogeneity of a unicellular population but also the near-unicellularity of the FLO1-expressing cells at low culture density. However, the strains had different absolute GFP/RFP ratios at the same oxygen level. This resulted from differences in individual fluorophore expression between the two strains (Figure S5D). Nonetheless, though the GFP/RFP ratio could not be directly compared between strains, within a given strain, it was a reliable readout of oxygen levels. We therefore focused on comparing DFOR reporter levels solely within the synthetic flocs, using culture density to control aggregate size.

To measure intracellular oxygen under normal aerobic conditions, we measured reporter output in a shaking incubator open to the atmosphere. Reporter output overlapped with what was observed in atmospheric oxygen on the eVOLVER (Figure S5E). We next cultured the synthetic flocs with DFOR in a shaking incubator starting from either low or high OD. We hypothesized that if oxygen was unable to reach cells at the center of an aggregate, we would observe a subpopulation of cells with much higher GFP/RFP ratios that specifically emerged at high culture density. However, reporter output at very low density (i.e., very little or no aggregation) overlapped with the output measured in very dense culture (Figure 5E). This was true when growing on a fermentable carbon source (i.e., glucose) as well as non-fermentable (i.e., glycerol) carbon source, where cells consume more oxygen.

The apparent lack of diffusion limitation was surprising given that snowflake yeast, which form much smaller aggregates, have limited aerobic respiration when grown in glycerol. This suggests that a crucial feature ensuring oxygen delivery in synthetic flocs could be cell rearrangement, which is possible in flocs but not in snowflakes.^33^ Overall, this result is consistent with our other findings suggesting that mass transport limitations are not a large concern for flocs, which could be general to other groups of cells held together by reversible cell-cell adhesion.

### Synthetic flocs grow better, not worse, in dilute sugars

Diffusion limitations are not merely a feature of aggregate size, but depend also bulk concentrations. While nutrient flux can exceed consumption if the bulk concentration is high, carbon could become limiting if bulk concentrations were lower. Combined theoretical and experimental work in yeast colonies has shown that glucose transport can become limiting at concentrations much lower than in standard rich media,^16^ though the absence of stirring and liquid media distinguishes colonies from synthetic flocs.

We therefore tested if synthetic flocs grew worse than unicellular controls in low glucose (Figure 6A). However, contrary to our prediction, as the glucose concentration decreased, synthetic flocs grew better, not worse, than the control mScarlet strain (Figure 6B). Moreover, this effect was larger for lower initial ODs (Figure 6C). This effect was not specific to glucose as synthetic flocs also grew better in limiting concentrations of other carbon sources, including other sugars (galactose, maltose) as well as non-fermentable carbon sources (ethanol, glycerol) (Figure 6D). Curiously, the advantage was absent in unstirred cultures, as all strains had similar endpoint ODs when grown in flat bottom plates that were only shaken for the first two hours of growth to allow aggregates to form (Figure S6A). While the reason for this remains unclear, we hypothesize that this could be because stirring is required for the collisions that generate flocs that otherwise flatten due to gravity, or because the unstirred culture becomes anaerobic more quickly.

**Figure 6.**
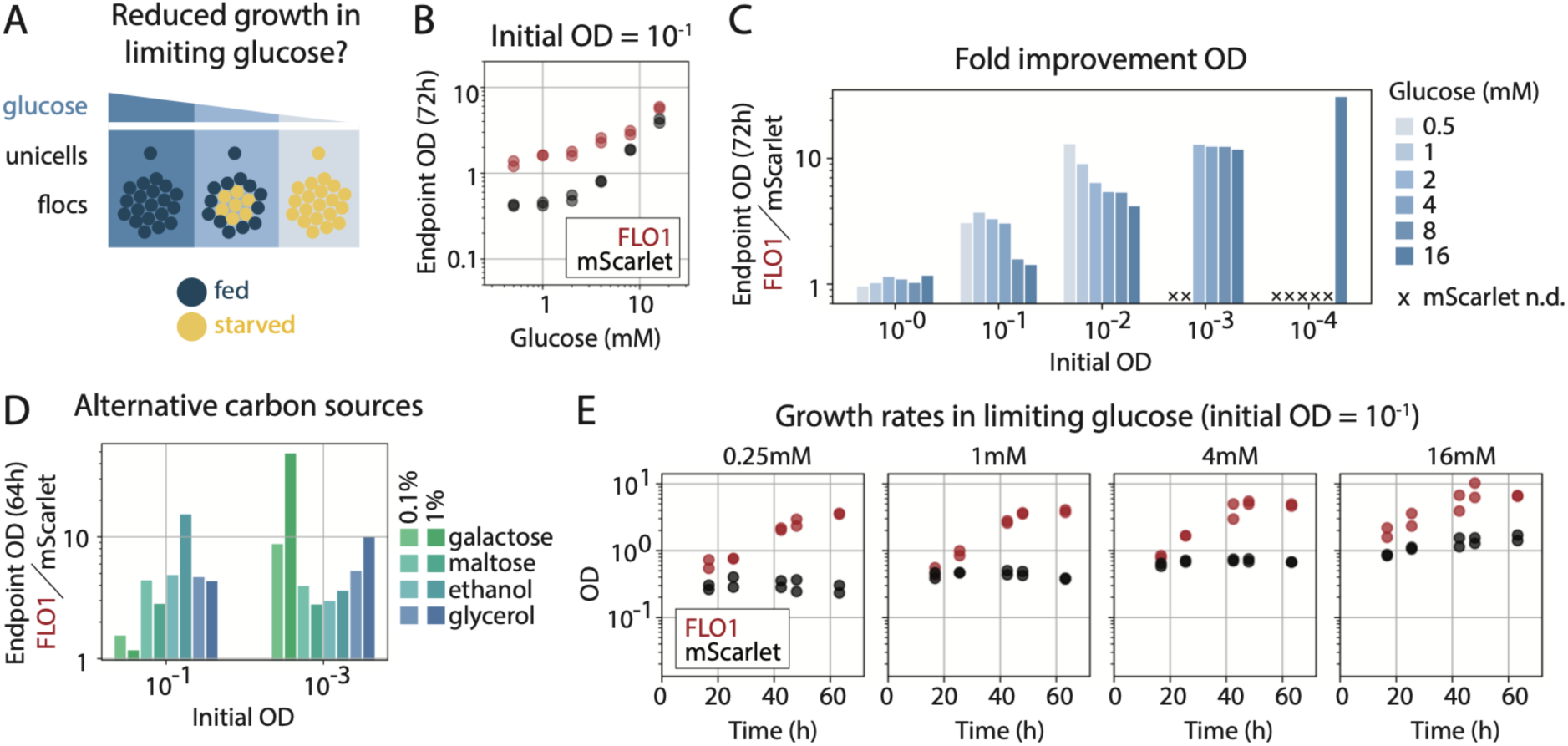
Aggregates grow better, not worse, in limiting carbon. **(A)** Diffusion limitations can increase when bulk concentrations are lower. While experiments in standard yeast media may contain sufficient glucose for transport to cells at the center of the aggregate, reducing the concentration of glucose may create conditions that starve central cells. **(B)** Inducible synthetic flocs and the mScarlet unicellular control were grown in YP media containing less than the standard ∼111mM glucose. Though both strains divide less frequently as the glucose concentration decreases, synthetic flocs significantly outgrow the mScarlet strain. **(C)** The higher endpoint OD of synthetic flocs relative to the mScarlet strain scales with the glucose concentration and initial cell density, with the synthetic flocs having the greatest advantage at low initial ODs and low glucose. In some cases, the mScarlet strain had no detectable growth, while synthetic flocs grew in all conditions. **(D)** Synthetic flocs also have higher endpoint OD when grown on YP media with alternative carbon sources, including a disaccharide (maltose) and non-hexoses (ethanol, glycerol). **(E)** Growth dynamics confirm that synthetic flocs have higher growth rates than the mScarlet strain on low glucose concentrations.

Synthetic flocs could have higher endpoint OD because of higher survival during an early “starvation” period after glucose was removed, rather than improved growth in low glucose. We therefore quantified early viability by plating countable colonies. However, in the absence of glucose, both strains had similar increases in viability at early timepoints (Figure S6B). In fact, the approximately 10-fold increase over 6 hours was consistent with the ∼1.7h doubling time we previously observed in rich media (cf. Figure 2B), suggesting that cells continue to divide using intracellular stores. Rather, differences emerged because of faster or merely sustained growth of the synthetic flocs, rather than loss of viability in those expressing mScarlet (Figure S6B). Measuring OD at later timepoints confirmed the faster or more sustained growth of the synthetic flocs produced their higher endpoint OD (Figure 6E). Similar trends were also observed in the constitutive variants of the synthetic flocs and mScarlet strain (Figure S6C).

Thus FLO1 expression, contrary to intuition, can improve growth when a carbon source becomes limiting. The higher benefits observed at lower sugar concentrations and lower culture densities are consistent with the effects of shared extracellular metabolites, which are more effectively captured in aggregates.^47^ As observed in our experiment, the benefits of metabolite sharing are largest when culture densities are low, producing larger distances between non-aggregating cells, and when metabolite concentrations are low.

### Benefits of non-specific flocculation can be privatized

Because non-specific FLO1 binding can incorporate strains with and without the adhesion protein in the same aggregate (Figure 7A), we hypothesized that its growth benefit in low carbon might also be non-specifically shared with other strains. We therefore mixed the inducible synthetic flocs and mScarlet strain at different ratios and measured growth in 1mM glucose, to see if the mixed aggregates grew better than unicellular strains in low glucose media (Figure 7B). All co-cultures of synthetic flocs and the unicellular strain grew better than the unicellular strain alone, and even co-cultures seeded with as few as 10% synthetic flocs grew as well as unmixed synthetic flocs (Figure 7C). However, the improved growth of mixtures was not due to cooperation between strains as we hypothesized. Using flow cytometry to check the proportions of the two strains over time, we found that the synthetic flocs outcompeted the non-aggregating strain (Figure 7D). While FLO1-expressing cells continued to grow exponentially, mScarlet strain growth saturated at the same lower OD in all mixtures (Figure S7A). Thus, the benefits of flocculation observed in low glucose were largely privatized to FLO1-expressing cells.

**Figure 7.**
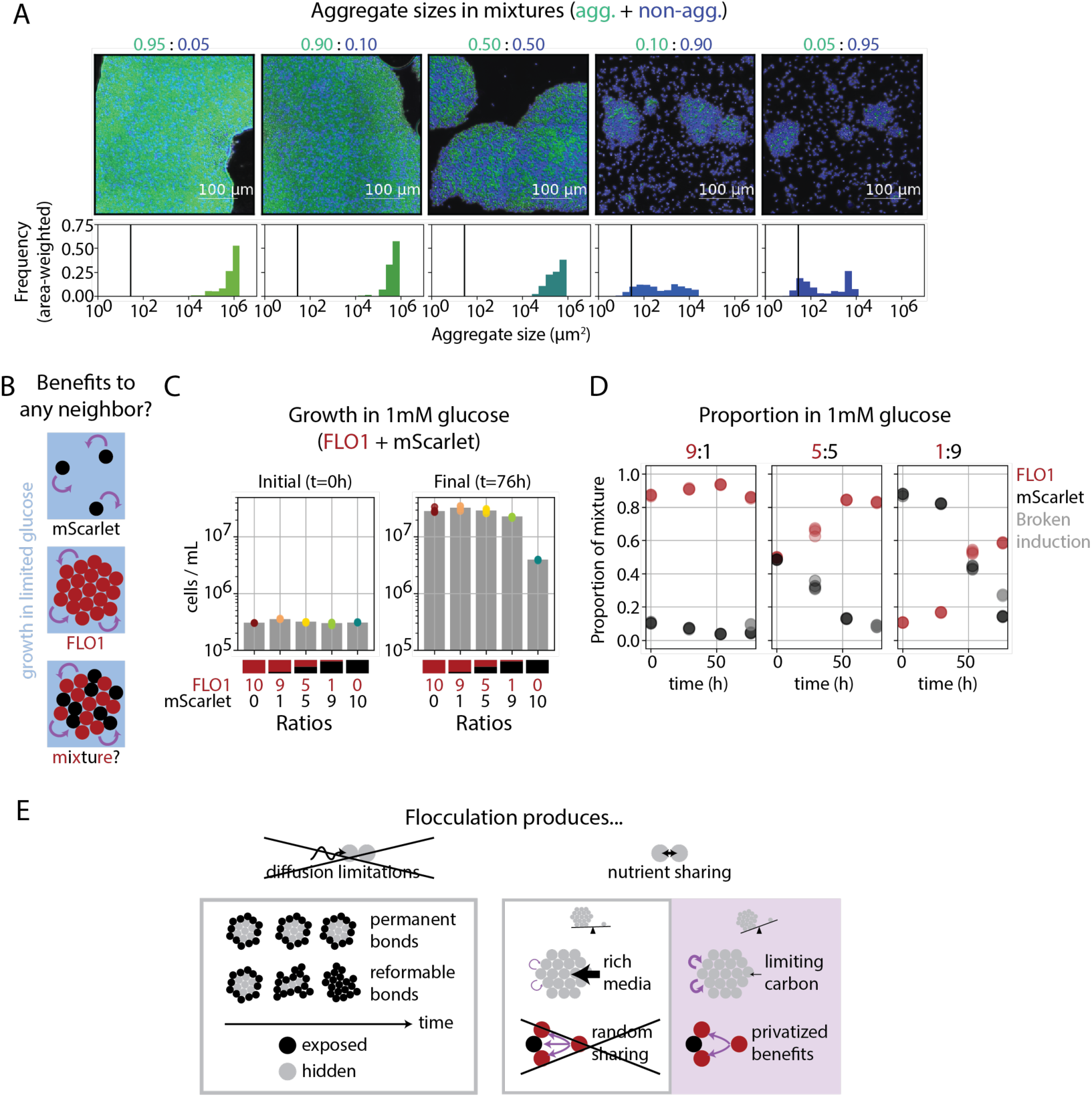
Though FLO1 binding is non-specific, FLO1-expressing cells selectively benefit in mixtures. **(A)** Constitutive synthetic flocs (green) were mixed at different ratios with a BFP unicellular strain. Aggregates can form even when very few FLO1-expressing cells are present, and cells need not express FLO1 to be incorporated into aggregates. However, aggregate size decreases when the proportion of synthetic flocs is below 90%. **(B)** We hypothesized that in low glucose, close proximity to other cells can improve growth, perhaps by sharing an extracellular metabolite. If true, then any neighbor could benefit from being present in an aggregate, whether or not they express FLO1. **(C)** Different mixtures of inducible synthetic flocs and the mScarlet strain were grown in 1mM glucose. Even mixtures containing only 10% synthetic flocs outgrew populations with the unmixed mScarlet strain. **(D)** Though mixtures start at very different ratios, synthetic flocs eventually outcompete the mScarlet strain. At the final timepoint, a third population with no mScarletI expression (i.e., much lower than the two competing strains) indicates a broken induction circuit. **(E)** Of the two possible effects of aggregation, flocculation does not produce diffusion limitations but can, in selective conditions, provide benefits of nutrient sharing. The reformable bonds of FLO1 can bring all cells in an aggregate in contact with the bulk environment. The benefits of nutrient sharing were only apparent in limiting carbon and, despite the non-specificity of binding, are selectively privatized to FLO1-expressing cells.

This privatized benefit was unexpected but could arise because aggregates are not randomly mixed. The “specific” interaction between cells expressing FLO1 could be stronger than the non-specific interaction between synthetic flocs and other genotypes.^48^ While flattening aggregates before imaging likely does not preserve spatial structure, we do see that only unicellular strains are excluded from aggregates when mixed (e.g., Figure 7A). Such a bias could further enrich FLO1-expressing cells at the aggregate center or in the largest aggregates. These larger aggregates could then reap proportionally larger benefits, as has been predicted in theoretical models^47^ and is consistent with our data. Constitutive synthetic flocs, which produce smaller aggregates, grew less well in low glucose than the larger induced synthetic flocs, though still better than both unicellular controls (Figure S7B). In this way, a “rich get richer” mechanism that enriches FLO1-expressing cells in the largest aggregates could allow them to outcompete other strains that only non-specifically enter aggregates.

## DISCUSSION

Engineering cell aggregates such that the benefits outweigh the costs is necessary for synthetic biology, but the same diffusion limitations and close proximity to neighbors that enable novel multicellular functions can also disrupt core cellular processes (Figure 1A). How to balance these tradeoffs is not obvious from studying extant multicellular organisms, due to their distant divergence from unicellular ancestors or dissimilarity from the most common, unicellular chassis strains. Thus, engineering aggregation in *S. cerevisiae* allowed precise comparisons between closely related “multicellular” and “unicellular” strains while also providing important insights into the effects of aggregation on a common synthetic biology chassis.

We produced synthetic flocs by engineering the expression of a single yeast adhesion gene, FLO1. High FLO1 expression generated aggregates containing 1000s of cells. However, such macroscopic aggregates had no detectable changes in growth rate and were stably maintained over 450 generations of continuous culture in rich media. RNA-Seq confirmed that expressing FLO1 had very limited, if any, effects on gene expression and suggested that previously identified differentially expressed genes may instead be side effects of high density, saturated cultures that produce the largest aggregates. FLO1 expression also did not provide protection from diverse chemical toxins, ranging from ethanol to a cell-wall degrading enzyme. Both the low fitness costs and limited physical protection were consistent with virtually absent diffusion limitations in our synthetic flocs, which could be fully stained by a 40kDa dye within 1 minute and a 100kDa dye within 1 hour. Though oxygen diffusion limitations were previously observed on a very short length scale (i.e., 3 cell diameters) in snowflake yeast, all cells in large flocs had similar levels of intracellular oxygen, suggesting cell rearrangement as a potential mechanism to boost nutrient access. Unexpectedly, decreasing sugar concentrations in an attempt to increase diffusion limitations actually improved aggregate growth relative to non-aggregating cells. However, this benefit was not equally shared among all cells in an aggregate. When mixed with a unicellular strain in limited glucose, FLO1-expressing cells overtook the culture. Overall, this suggests that the benefits of flocculation are mostly related to extracellular metabolite sharing, rather than protective diffusion limitation, and could be specific to nutrient-limited settings (Figure 7E).

These findings in synthetic flocs suggest possible design principles that could be applied to engineering multicellularity. First, reversible cell-cell bonds may limit fitness costs associated with diffusion by allowing cell rearrangements. Intentional use of reformable bonds could increase mass transport into mammalian cell aggregates. By contrast, when diffusion limitations are desired for protection, synthetic adhesins that form covalent bonds, or unstirred culture conditions where cells cannot exchange places, could be valuable. FLO1 may also permit diffusion because it is not as space filling as extracellular matrix, which is present in many biofilms, organoids, or even some wildtype flocs.^22^ Constructing such an ECM with polysaccharides,^2^ secreted adhesion proteins,^49^ or even hydrogels provides an alternative mechanism to increase protection in yeast aggregates. Second, beyond access to molecules in the bulk, cells in aggregates have higher access to cell-derived molecules, which we hypothesize as the mechanism of their improved growth in limiting carbon. Consistent with this, higher metabolite exchange was recently shown in synthetic flocs, both for crossfeeding strains and two strains with different halves of a biosynthetic pathway.^5^ The robustness we found in low nutrient settings further suggests that engineered aggregation could be valuable for bioremediation strains or biomanufacturing, where cells do not have constant access to rich media.

FLO1’s ability to produce larger aggregates than synthetic adhesins also suggests potential strategies for engineering novel adhesion proteins. FLO1 and FLO10 have similar mannose-binding domains, but FLO10 does not produce aggregates (Figure 1C). Similarly, the same synthetic adhesins produce larger aggregates when displayed on stalks rather than as Aga2 fusions. This echoes prior studies of stalk variation in FLO proteins^22,25,26^ where domains outside the adhesin domain tune overall avidity of two strains for each other. This suggests that we can generate additional diversity in adhesion strength without mutating the binding interface and that screening naturally occurring variation in FLO proteins could generate a toolkit for display parts. Similarly, the large sizes of FLO1-based aggregates suggest that weak binding of abundant cell-surface moieties can be a useful mechanism for producing aggregates with tunable sizes, rather than developing extremely high affinity binding partners, as they access benefits of multivalency without requiring high expression of the ectopic construct. FLO1 belongs to a class of sugar-binding cell surface proteins called lectins that are also prevalent in the plant and animal kingdoms.^50^ Though these lectins largely serve important immune functions identifying foreign glycosylation patterns, displaying sugar-binding domains on synthetic stalks could provide synthetic adhesion proteins with strong multivalent binding in mammalian cells.

FLO1 is also an attractive platform for living materials as its limited fitness effects, as well as its homotypic and non-specific binding, allow it to synergize with existing engineering approaches. For example, any cell type expressing FLO1 can be flexibly paired with another yeast strain, even at very low abundance (cf. Figure 7A), to seed aggregation in the co-cultured strain. In another approach, strains engineered to rapidly convert a toxin, rather than passively absorb it, could generate the necessary diffusion limitations to protect an inner core of cells. This has analogies with biofilms that can generate protective diffusion limitations by expressing catalase to break down hydrogen peroxide.^51^ FLO1 expression could also synergize with other co-expressed adhesion molecules, such as with SpyTag and SpyCatcher, to make attachments longer-lived. Alternatively, co-expression with proteins for surface attachment could seed large, surface-associated aggregates. This approach could facilitate the breakdown of macroscopic substrates like recycled plastic, where flocculation produces locally high concentrations of extracellular enzymes while the other adhesin localizes the cells to their target.

Identifying costs and benefits associated with synthetic cell-cell adhesion also has implications for the evolution of multicellularity. Non-specific binding that allows multiple genotypes to co-exist in the same aggregate allows “cheater” cells to benefit from aggregation without investing energy into aggregate formation, essentially breaking natural selection for aggregation.^17^ However, our synthetic flocs could outcompete, though not eradicate, a co-cultured unicellular strain that was non-specifically incorporated into aggregates. This could arise from a bias toward specific interactions over non-specific ones, where FLO1-expressing cells are enriched in aggregates relative to their unicellular competitors.^22^ Such a co-existence strategy could provide an evolutionary route to more robust multicellularity, if limiting the growth of the competing strain buys time for the non-specific binder to evolve cheater-excluding specificity. A similar effect has been recently observed in experimental evolution of flocs.^52^

While we identified a culture context (i.e., low sugar) that significantly alters the relative fitness of aggregating and non-aggregating strains, there are likely many other contexts that produce different and larger differences between aggregating and non-aggregating fitness. These could span growth conditions, adhesion mechanisms, and genotype, as our synthetic flocs did not recapitulate all behaviors of wildtype flocs. Studying wildtype flocs could reveal which genes modulate the positive and negative effects of flocculation,^53^ as preexisting traits could facilitate the persistence of aggregation or aggregation can mask the effects of otherwise costly traits.^9^ Large scale studies of growth and fitness of different strains expressing FLO1 could identify which cellular processes might fit into these categories. Such relationships between cell aggregation and core cellular processes will likely comprise the next generation of design principles for engineering synthetic multicellularity.

## AUTHOR CONTRIBUTIONS

H.E.K., A.S.K., and M.J.D. conceived and designed the study, with input from S.L., K.T., and D.H. H.E.K., S.L., and K.T. engineered yeast strains. H.E.K. and S.L. performed the longterm evolution experiment in base eVOLVER, and H.E.K. and D.H. measured DFOR in the atmospheric control eVOLVER. H.E.K. implemented the remainder of the experiments and conducted data analysis. H.E.K., A.S.K., and M.J.D. wrote and edited the manuscript. All authors reviewed the manuscript and approved the final version of the manuscript for submission.

## ACKNOWLEDGEMENTS

We thank members of the Dunlop and Khalil labs for input. We thank Mikel Lavilla-Puerta and Francesco Licausi for sharing the DFOR construct. We also thank William Shaw and Tom Ellis for sharing constructs for generating the base strain used in this study as well as parts for synthetic adhesins. This work was supported by NIH grants R01EB027793 and R01AI171100, Schmidt Sciences Polymath award G-22-63292, and the Vannevar Bush Faculty Fellowship (VBFF) under Grant N00014-20-1-2825. H.E.K. was supported by Boston University’s Kilachand Fellowship and a Damon Runyon Cancer Research Foundation (DRG-2472-22). M.J.D. acknowledges support from the National Science Foundation through grant MCB-2143289.

## RESOURCE AVAILABILITY

### Lead Contact

Further information and requests for resources and reagents should be directed to and will be fulfilled by the Lead Contact: Ahmad S. Khalil

### Materials Availability

Key plasmids will be deposited at Addgene for distribution. DNA constructs and strains are available from the lead contact.

### Data and Code Availability

- Raw RNA-seq data for transcriptome analysis will be deposited in the NCBI GEO database. Accession number will be listed in key resources table. All other raw datasets will be deposited on Dryad. Associated DOI will be listed in the key resources table.
- All original code will be available on Github and will be deposited on Zenodo. Associated DOIs will be listed in the key resources table.
- Any additional information required to reanalyze the data reported in this paper is available from the Lead Contact upon request.

## MATERIALS AND METHODS

### Bacterial strains and growth media

*E. coli* strains DH5α and TOP10 were used for all cloning experiments. Selection and growth of *E. coli* were performed in Lysogeny Broth (LB) medium at 37°C with aeration. When necessary, the LB medium was supplemented with the appropriate antibiotics for plasmid maintenance (chloramphenicol 34μg/mL, carbenicillin 100μg/mL, kanamycin 50μg/mL, or spectinomycin 100μg/mL). See Supplemental Table S1 for a list of plasmids used in this study.

### Bacterial transformations

Chemically competent *E. coli* cells were created following the TSS protocol for KCM transformations.^54^ A colony of *E. coli* was grown to saturation overnight in 10 mL of LB and then split into two 2 L baffled flasks with 500 mL of LB. The culture was grown for 2–3 h to OD600 ∼ 1.0, chilled on ice to stop growth, and centrifuged at 4000*g* at 4°C for 10 min. The supernatant was then discarded, and the cell pellets were resuspended by aspiration in 100 mL of ice-cold TSS (85 mL of LB, 10 g of PEG-3350, 5 mL of DMSO, and 2 mL of 1 M MgCl_2_). 100μL of the cell suspension was then aliquoted into 0.2 mL PCR strip tubes, flash frozen in liquid nitrogen (or dry ice), and put into a −80°C freezer for long-term storage. To transform the DNA, 100μL of 2× KCM (200 mM KCl, 60 mM CaCl2, 100 mM MgCl2) was added to 100μL of the competent cells after 10 min of thawing on ice. 20µL of the competent cell-KCM cocktail was then added to 1–4µL of DNA and kept on ice for 5 min. The cells were then heat-shocked in a water bath at 42°C for 1 min, transferred back to ice for 1 min, and then recovered at 37°C for 1 h (shaking not necessary). Cells were then plated on solid LB medium supplemented with the appropriate antibiotics.

### Yeast strains and growth media

*Saccharomyces cerevisiae* strain BY4742 (*MAT*α *his3*Δ*1 leu2*Δ*0 lys2*Δ*0 ura3*Δ*0*) was used for all experiments. Yeast extract peptone dextrose (YPD) media was prepared with 1% (w/v) Bacto Yeast Extract (Merck), 2% (w/v) Bacto Peptone (Merck), 2% (w/v) glucose (VWR). Synthetic Complete (SC) medium was prepared with 2% (w/v) glucose (VWR), 0.67% (w/v) Yeast Nitrogen Base without amino acids (Sunrise Science Products), and 0.8 g/L complete supplement mix (CSM; Sunrise Science Products). For auxotrophic selection, SC medium minus the appropriate supplement was used instead of CSM (SC-Ura, SC-Leu, and SC-His; Sunrise Science Products). For NAT selection, 100µg/mL nourseothricin was added to the SC media. Solid medium was prepared with 2% (w/v) agar (VWR). Inducible strains were first grown overnight in YPD, backdiluted in YPD to an OD of 0.2 for 4-8h of growth, and then backdiluted to an OD of 0.05 in YPD with aldosterone for a second overnight culture. Cultures were then maintained at the same aldosterone concentration throughout the experiment. Strains were induced with 5μM aldosterone unless otherwise specified. Yeast strains were generated by transformation with NotI-digested plasmids, as described below. See Supplemental Table S2 for a list of all strains used.

### Yeast transformations

Yeast cells were transformed using the lithium acetate protocol, adapted from Gietz and Woods.^55^ A yeast colony was picked from a plate and grown to saturation overnight in 2 mL of YPD medium. The following morning the cells were diluted 1:100 in YPD (OD600 ∼ 0.175) and grown for 4–6 h to OD600 0.6–0.8 (1-3mL of culture corresponds to a single transformation). Cells were pelleted at 3000*g* for 3 min in a large benchtop centrifuge at room temperature and washed once with half the starting volume of 0.1 M lithium acetate (LiOAc) (Sigma) and repelleted. Cells were then resuspended in 0.1 M LiOAc to a total volume of 100µL/transformation. 100µL of cell suspension was then distributed into 1.5 mL reaction tubes and pelleted at 8000 rpm for 3 min on a small benchtop centrifuge at room temperature. Cells were resuspended in 64µL of DNA-salmon sperm DNA mixture (10µL of boiled salmon sperm DNA (Invitrogen) + DNA + ddH2O, where DNA was prepared as below) and left to incubate at room temperature for 10-30 min. 294µL of PEG-LiOAc mixture (260µL 50% (w/v) PEG-3350 (Sigma) + 36µL 1 M LiOAc) was then added, gently vortexed for five seconds, and incubated at room temperature for 30 min. The yeast transformation mixture was then transferred to a water bath at 42°C for 15 min. Cells were pelleted at 8000 rpm in a small benchtop centrifuge for 20 s at room temperature and resuspended in 0.1–1 mL of sterile water (for auxothrophic selection) or YPD (for antibiotic selection), and plated onto the appropriate synthetic dropout medium (after 10 min at room temperature) or antibiotic medium (after 3 h recovery at 30°C).

DNA for transformation was digested with NotI as follows: plasmid DNA (various amounts), 1µL of CutSmart buffer (NEB), 0.2µL of NotI (NotI-HF, NEB), and up to 10µL of sterile H2O. DNA was digested at 37°C for 4h or overnight (∼16h). The entire reaction was then combined with water and boiled salmon sperm DNA to 64μL, as described above, and directly transformed into yeast without a cleanup step. For multiplexed markerless CRISPR-Cas9 experiments using more than four integration vectors, integration vectors should be individually digested and column purified before transformation to achieve high efficiency.

### OD and growth rate measurements

#### All OD measurements are OD_600_

Aggregates must be broken up into single cells to reliably measure OD by spectrophotometer. Because calcium ions are required for flocculation, EDTA, which chelates divalent cations, can be used to create a single-cell suspension. To measure OD, cultures were diluted at least 1:2 in 100mM EDTA, for a final concentration of 50mM EDTA. However, for very high FLO1 expression, dilution in 500mM was required to break up aggregates. Often, further 3-fold serial dilutions in 100mM EDTA were performed to reach an OD between 0.1 and 0.4, the reliable linear range of our spectrophotometer. In parallel, an identical volume of growth media was similarly diluted in EDTA for background subtraction at each dilution. OD was then measured by subtracting the appropriate background and multiplying by the dilution. In some cases, multiple raw OD measurements from this serial dilution curve fell within the desired linear range. In this case, we took the median of all these values (after background subtraction and dilution correction) as the best estimate of the sample OD.

To measure growth rates, the same starting culture was partitioned across replicate wells in deep 96-well blocks (500μL per well with 1.2mL fill volume) and a number of blocks equal to the number of timepoints. All blocks grew in parallel in the shaker. At the specified timepoint, a block was removed, and the OD measured in all wells as described above. To infer growth rate, the log of each strain’s OD over time was fit by least squares to a linear model, with at least three replicate wells measured per timepoint.

### Size quantification by microscopy

To image aggregates, cultures were backdiluted to the same OD (see “OD and growth rate measurements”) in 24 or 96 well blocks and returned to the shaking incubator (37°C) for 1-3h hours to allow aggregates to reform. The initial OD and time for aggregate reformation was modified to suit the experiment. For example, larger aggregates should start from lower ODs to limit aggregate sizes, and growing for shorter intervals limits loss of aggregates to the walls of the wells. Nonetheless, any time sample sizes were to be compared, all samples were prepared at the same initial OD and backdilution period, often cultured on the same block.

After aggregates reformed, 2-3μL of culture was sampled with cut-off P20 tips (whose larger inner diameter limited aggregate break-up) as quickly as possible, as aggregates could settle and fuse as blocks sat without stirring, and dropped on a glass slide. In some instances, glass slides were pre-treated with ConA, which binds the cell wall to hold cells in place, by spreading a thin layer of 1 mg/mL solution over the slide and letting it dry over 15 minutes.^9^ Quickly after plating, a glass coverslip was carefully dropped from a few millimeters above the culture droplet, starting from a slight angle relative to the glass slide to avoid trapping air bubbles as it fell. When imaging many conditions at a time, samples were plated in batches, with the block being returned to the shaker in batches. Slides were then imaged with a Nikon Eclipse Ti2, capturing multiple fields of view per sample.

To quantify aggregate sizes, cells were segmented on a constitutive fluorescent reporter using packages from scikit-image. Images were binarized using Otsu’s method to automatically threshold pixel values. Aggregates were identified using binary closing with a 5 pixel disk to close holes. With an interpixel distance of 0.33μm for images acquired at 20x magnification, this corresponds to a distance of ∼1.5μm, the approximate maximum distance between cells that were in the same aggregate. The sizes of the resulting objects in pixels were converted to the aggregate area by multiplying by the interpixel distance squared. For aggregates composed of mixed strains expressing different segmentation colors, we thresholded each channel individually and summed their boolean masks before binary closing.

To visualize the distribution of aggregate sizes, we calculated the area-weighted probability. While we might consider the probability of observing an aggregate equal to 1 divided by the number of aggregates, this underrepresents the number of large aggregates. For example, a sample with only a 100-cell aggregate and a 2-cell aggregate is not 50% 100-cell aggregates. Instead, we preferred a measure closer to the probability of an individual cell being incorporated into an aggregate of a particular size. Therefore, we instead calculate the probability weighted by the area. Each aggregate’s size is divided by the sum of all aggregate sizes observed for that sample, usually across more than one field of view. To plot the histogram of this quantity, we calculate the sum probability of all aggregates within a particular bin (i.e., range of sizes).

### Viability measurements

To estimate viability, as with estimating OD, aggregates must be broken into a single-cell suspension prior to plating countable colonies. However, no colonies grew from cultures diluted in 50mM EDTA, suggesting EDTA was toxic at those concentrations. Therefore, we first diluted aggregates 1:3 in 100mM EDTA, but then diluted that single-cell suspension in PBS to reach a final 200-fold dilution. This solution was then serially diluted 4-fold seven more times. 6μL of each dilution was then plated on a YPD agar plate. This sets our lower limit of detection around 100,000 cells/mL (approximately OD 0.01), as a 6μL droplet of the lowest dilution has a 95% chance of containing at least one cell.

Plates were scanned and colonies manually counted two days after plating. Wells containing approximately 30 colonies or fewer (including zero) were included for analysis. We assumed that colony counts followed a Poisson distribution, where the concentration of viable cells was the expected number of events. We estimated this rate for each sample by performing maximum likelihood estimation on all observations of countable colonies and the associated dilutions for that sample. This rate was estimated independently for technical replicates, i.e., wells that were independently grown, diluted, and plated.

### Continuous culture for 450 generations

To continuously culture aggregates, we used the eVOLVER platform.^34^ A significant challenge was running the experiment in rich media (i.e., YPD) without experiencing contamination. Therefore, we prepared 20L media carboys by autoclaving 2L of water with a 45min liquid sterilization cycle, before adding 20L of media, and sterilizing for 3 hours. For further protection against bacterial contamination, we added chloramphenicol (final concentration 30μg/mL) and carbenicillin (final concentration 50μg/mL) to the media. We also used rubber stoppers to plug the sample port of the eVOLVER lids. Lastly, we loaded rich media into all the eVOLVER vials and waited five days to ensure nothing was growing before inoculating our yeast strains.

Cells were cultured in turbidostat mode, set to maintain the target OD between 0.65 and 0.75, though we note that this measurement is less accurate for aggregates which scatter light differently than a single cell suspension. This OD range produced large, visible aggregates for high FLO1 expression and was near the maximum that can be measured on eVOLVER. The narrow OD range ensured that aggregate size did not change much throughout the experiment and kept the cells constantly in exponential phase, maximizing the number of generations reached over the 29 days of the experiment. On the final day, strains were sampled from eVOLVER and imaged alongside fresh overnight cultures of the same strain. To quantify sizes, all samples were backdiluted to an OD of 0.8 (see “Size quantification by microscopy” for full protocol). Before flow cytometry, cells were diluted 1:3 in 500mM EDTA to produce a single-cell suspension. Endpoint samples for “evolved” strains were also frozen in 25% glycerol at -80°C. These samples were thawed after the fact for ploidy analysis (see “DNA ploidy staining”).

### Growth in toxic conditions

For all toxic treatments, cells were induced with 5μM aldosterone prior to the start of the experiment (see “Yeast strains and growth media”). Cultures were then backdiluted to approximately 80% of the desired starting OD, i.e., 0.8 and 4 to reach 1 and 5, respectively. Each culture, representing one induced strain at one of the two starting densities, was split between multiple wells such that at each timepoint, for each concentration of toxin, OD could be measured in triplicate. Wells for the same timepoint were all put on the same block, such that one block was sacrificially measured at each timepoint.

Blocks grew for 2h at 37°C in a shaking incubator to allow aggregates to reform. At the start of the experiment, at least 6x concentrated toxin or an equivalent volume of water was added carefully and quickly to the side of each well. All blocks except one were returned to the shaker for mixing. This ensured that aggregate size was not perturbed by pipetting aggregates up and down or drastically changing the volume of the culture.

The leftover plate was measured as the first timepoint, including measuring OD (see “OD and growth rate measurements”) and plating for measuring viability (see “Viability measurements”). OD measurements, which could be completed more quickly, were taken approximately every thirty minutes, while countable colonies were plated at the start, middle, and end of the experiment.

For ethanol and amphotericin, toxin was added to YPD for measurements of growth and viability. However, we found that Zymolyase’s activity was much reduced in YPD. Therefore, Zymolyase was diluted in HBSS, with supplemental calcium to ensure that flocs could still form. While cells stayed alive in the buffer, they did not continue to grow and divide. In this more aqueous media, flocs had a higher propensity to stick to the walls of the wells, creating an apparent decrease in OD. Therefore, we measured OD over shorter time scales, both because Zymolyase was still active in that period and to limit loss to the walls.

### RNA sequencing (RNA-seq)

To prepare samples with and without flocculation at different culture densities, non-aggregating (yHK74) and aggregating (yHK66) strains were grown overnight in rich media and then backdiluted 1:200 the following morning. The day after, these cultures were split and diluted to an OD of either 0.25 or 2.5. Six replicate wells were plated across different 24well blocks and returned to the shaking incubator. After 4h, one plate was removed and imaged to check aggregate sizes (see “Size quantification by microscopy”). A second plate was used to measure OD, which confirmed ODs of 1.2 and 1.3 for the dilute condition and 7.2 and 7.0 for the dense condition (for yHK74 and yHK66, respectively). The remaining block, with three replicates, was then removed from the shaking incubator and put on ice. 1.5mL of the dilute condition and 0.3mL of the dense condition were taken, to ensure a similar number of cells in all samples. These cultures were then pelleted.

RNA was isolated from these pellets with Zymo’s Direct-zol RNA Microprep kit. Pellets were resuspended in 800μL TRI reagent in a chemical fume hood and transferred to RNAse free tubes with glass beads. These tubes were loaded into Qiagen’s Tissue Homogenizer, which was shaken at max speed for two cycles of 60s. This lysate was then purified on the Zymo column per manufacturer’s instructions, including an on-column DNAse digestion. Equal masses of purified RNA were used to create cDNA (Bio-Rad, iScript cDNA Synthesis kit) from each replicate.

Sequencing libraries were prepared from these samples with NEB’s UltraExpress mRNA kit. Libraries were then sequenced on Illumina NextSeq 2000 with a P2 flow cell, whose 400M reads ensure >30M reads per sample. We collected paired-end reads over 200 cycles. Read quality was assessed with FastQC, followed by genome alignment with STAR. Pseudo-counts were then generated with Salmon and checked by Qualimap. From this count matrix, we identified differentially expressed genes with PyDeSeq2. We fit a linear model on the effects of FLO1 expression and cell density. We identified differentially expressed genes considering both a cut-off of both significance (log_10_ adjusted p-value < 0.05) and effect size (fold-change > 1.5).

### Dyeing aggregates

For dyeing experiments, single colonies were picked into overnight cultures, diluted 1:200 the following day, and backdiluted to OD ∼1 in HBSS supplemented with calcium the day after that. These were split between 3 blocks, with one block for each technical replicate. Each technical replicate included two wells of each strain, one for dyeing and one control for background dyeing. Plates were returned to the shaking incubator to allow aggregates to reform. The blocks were processed in series after 1h (the first) or 2h (the third) of shaking.

On each block, half of the wells received the dye, while the other half did not. For wells that received the dye, the dye was added as a 1:5 dilution (i.e., 100μL dye to 400μL culture) against the wall of the well, to limit the volume changes that change aggregate size and large flows that could break up aggregates. Dyes were added by multi-channel pipet, so all strains received the dye at the same moment. Dyes were added at a final concentration of 0.05ug/mL Calcofluor White (Biotium, 29067), 30ug/mL Wheat Germ Agglutinin CF 405S (Biotium, 29027), and 100ug/mL concanavalin-A CF 405S (Biotium, 29075). The plate was quickly returned to the shaker and shook for 80 seconds. Then, the dyeing reaction was “quenched” by quickly adding 9mL of YPD, which reduced the concentration of the dye nearly 20-fold. For the “unstained” controls on each plate, YPD was first mixed with an equivalent 100μL volume of dye before being added to each well. This captures how much the cells are dyed in the presence of dilute dye alone during the centrifugation steps. We tried dilution with PBS, but found that it reversed some of the on-target staining and did not remove background staining as effectively.

To remove the dye and break up the aggregates, cultures were spun down (3000g, 3min), washed in 9mL PBS, spun again (3000g, 3min), and finally resuspended in 1mL of PBS. Note that the absence of calcium in PBS ensures that aggregates cannot form. Cells were then analyzed on the Attune NxT.

### Intracellular oxygen measurements with DFOR and Atmostat eVOLVER

Intracellular oxygen was quantified with the Double Fluorescence Oxygen Reporter, a fluorescent variant of the Double Luciferase Oxygen Reporter^46^ generously provided by Mikel Lavilla Puerta. This reporter was integrated into “dark” versions of the unicellular and flocculating strains, expressing mutFLO1 and FLO1 respectively. To verify the performance of the reporter at different oxygen conditions, we grew both DFOR reporter strains at low culture OD (i.e., turbidostat with target OD of 0.3) under atmostat control. The atmostat is a specially designed eVOLVER module that maintains oxygen levels at a desired setpoint. Because constant bubbling is required for efficient gas transfer, Antifoam 204 was added to culture media to limit foaming. The 16 vials were used to hold two replicates of each strain at 0, 2, 5, or 20% oxygen. These concentrations were regulated by bubbling a mixture of pure industrial grade nitrogen gas (N_2_) and air, corresponding to 100% N_2,_ 90% N_2_ + 10% air, 75% N_2_ + 25% air, and 100% air, respectively. After 24h of growth with constant dilution to maintain the low OD, 200μL was sampled from each vial and added to a solution of EDTA (100mM final) and cycloheximide (Sigma #01810, 20µg/mL), to break up aggregates and inhibit translation, to ensure the fluorescence of the cells was not affected by contact with the aerobic conditions outside the vial. These samples were then analyzed by flow cytometry (Attune NxT).

To assess intracellular oxygen inside flocs grown in aerobic conditions, the flocculating reporter strain was grown overnight in YP with 2% glucose (i.e, YPD) or YP with 2% glycerol. The following morning, the two cultures were serially diluted on the same carbon source to ODs ranging from 1 to 0.01, where the very low density showed the behavior of the reporter when aggregates were not formed. Two replicates of each were plated in a 96well block and grown for four hours in the shaking incubator. 200μL of culture was again disaggregated with EDTA and translation inhibited with cycloheximide.

### Growth in low glucose

Similar to other growth assays, strains were pre-grown in YPD before the start of the experiment. Colonies were picked into YPD on the first day, and overnight cultures were diluted into larger volumes on the second day to provide enough cells for the start of the experiment on the third day. If strains were inducible, induction was started on the second day.

To remove sugar, cultures were spun down on the third day and resuspended in YP containing no sugar. After measuring OD, cultures were normalized to an OD of 1 and then serially diluted 10 fold. To mix aggregating and non-aggregating genotypes, we further mixed these cultures at known ratios, such that the total cell density was the same. To these cultures, distributed between replicate wells on 96 well plate blocks, sugar (prepared in water at 20x the desired final concentration) was added at a 1:20 dilution. Unstirred conditions were initially stirred for 2h in the shaking incubator to allow aggregates to form, before being moved to a 37°C incubator with no shaking. Stirred conditions were incubated in the shaking incubator for the duration of the experiment. At the indicated timepoint, OD was sacrificially measured in each well as described above (“OD and growth rate measurements”).

To quantify OD at earlier timepoints when cell density was below the limit of detection for our spectrophotometer, we either plated countable colonies as described above (“Viability measurements”) or inferred cell density from our flow cytometry data, using the number of events as the number of cells in the (known) sample volume. To quantify what fraction of cells were aggregate versus not aggregating, we gated these events on RFP, which was higher in the non-aggregating strain that contained two copies of the mScarlet-I gene. We identified a population of cells with a broken induction circuit that had lower RFP levels than either strain, and therefore could originate from either population.

### DNA ploidy staining

To measure ploidy after the long-term culturing in YPD media, we followed a published protocol for ploidy staining.^56^ Briefly, we thawed glycerol stocks of each vial stored immediately after the long-term culturing experiment, alongside glycerol stocks of the parental strains. To preserve possible population diversity, we directly inoculated a large frozen volume of the glycerol stock into YPD. These were passaged 1:100 in YPD, and then stained the following day.

Before staining, overnight cultures were backdiluted 1:20 to allow 2 hours of exponential growth. These cultures were then diluted to an OD of 0.6 in 1.5mL before being spun down (500g, 3min) and resuspended in an equal volume of 70% ethanol. After a 90 minute fixation, cells were spun down and resuspended in 50mM sodium citrate twice. To reach single cells, this suspension was sonicated at 30% power for 30s. This sample was again spun down and resuspended in 200μL of 50mM sodium citrate with 0.5mg/mL RNAse. After a 2.5h incubation at 37°C, the volume was split between a 15μL unstained control and 185μL cell suspension for staining. To this was added 5μL of 1mg/mL propidium iodide and 5μL of 50mM sodium citrate. This sample incubated almost 24h in the dark at 37°C and then stored at 4°C for 3 days. To prepare samples for flow, the cell suspension was sonicated again, and 5μL of the stained sample was diluted 1:40 in 25ug/mL propidium iodide in 50mM sodium citrate.

