## Supplemental Information for "Synthetic cell-cell adhesion provides benefits of proximity without diffusion-related costs"

### SUPPLEMENTAL FIGURES

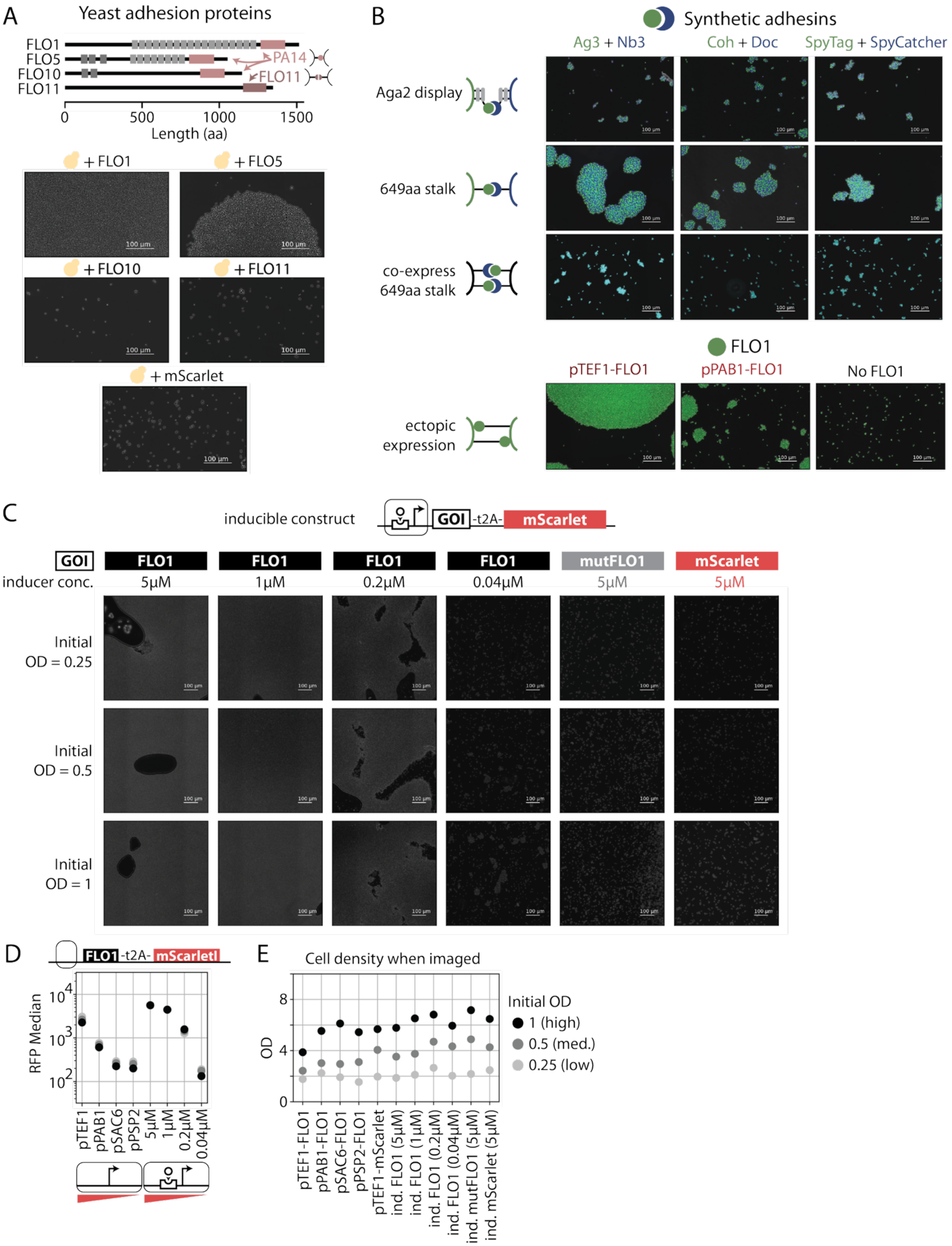
A
Yeast adhesion proteins
FLO1
FLO5
FLO10
FLO11
PA14
FLO11
0
500
1000
1500
Length (aa)
+ FLO1
+ FLO5
+ FLO10
+ FLO11
+ mScarlet
100 µm
B
Synthetic adhesins
Ag3 + Nb3
Coh + Doc
SpyTag + SpyCatcher
Aga2 display
649aa stalk
co-express 649aa stalk
FLO1
pTEF1-FLO1
pPAB1-FLO1
No FLO1
ectopic expression
C
inducible construct
GOI
-t2A-
mScarlet
GOI
inducer conc.
FLO1
5µM
FLO1
1µM
FLO1
0.2µM
FLO1
0.04µM
mutFLO1
5µM
mScarlet
5µM
Initial OD = 0.25
Initial OD = 0.5
Initial OD = 1
D
FLO1
-t2A-
mScarlet
RFP Median
$10^4$
$10^3$
$10^2$
pTEF1
pPAB1
pSAC6
pPSP2
5µM
1µM
0.2µM
0.04µM
E
Cell density when imaged
OD
8
4
0
Initial OD
1 (high)
0.5 (med.)
0.25 (low)
pTEF1-FLO1
pPAB1-FLO1
pSAC6-FLO1
pPSP2-FLO1
pTEF1-mScarlet
ind. FLO1 (5µM)
ind. FLO1 (1µM)
ind. FLO1 (0.2µM)
ind. FLO1 (0.04µM)
ind. mutFLO1 (5µM)
ind. mScarlet (5µM)

**Figure S1. FLO1 provides best single-genotype control of aggregate size.**

(A) Yeast flocculation proteins are all homotypic, with either a non-specific (PA14) or specific (FLO11) adhesion domain. Stalks contain various repeats of flocculin (light gray) and flocculin type 3 (dark gray) domains, with annotations taken from UniProt. Despite these similarities, not all flocculation proteins produced aggregates. Microscopy images are aggregates from overnight cultures flattened between a coverslip and glass slide into a 2-D monolayer.

(B) Synthetic adhesins only produce large aggregates when displayed with a long, 649-amino acid stalk, but this effect is abolished when the two binding partners are expressed in the same cell. By contrast, FLO1 provides the simplicity of a single genotype, as well as a larger range of aggregate sizes.

(C) Controlling FLO1 expression with a small-molecule inducible promoter produces similar effects as tuning expression with constitutive promoter strength (cf. Figure 1). The mutFLO1 and mScarlet unicellular controls do not produce aggregates.

(D) Comparing co-transcribed mScarlet expression in the constitutive and inducible expression systems shows similar levels of FLO1 expression, with maximal induction achieving higher levels than the pTEF1 promoter.

(E) OD measurements of cultures prior to imaging show that the inducible systems (imaged in Figure S1C) were assayed at similar ODs as the constitutive ones (imaged in Figures 1C and 1D).

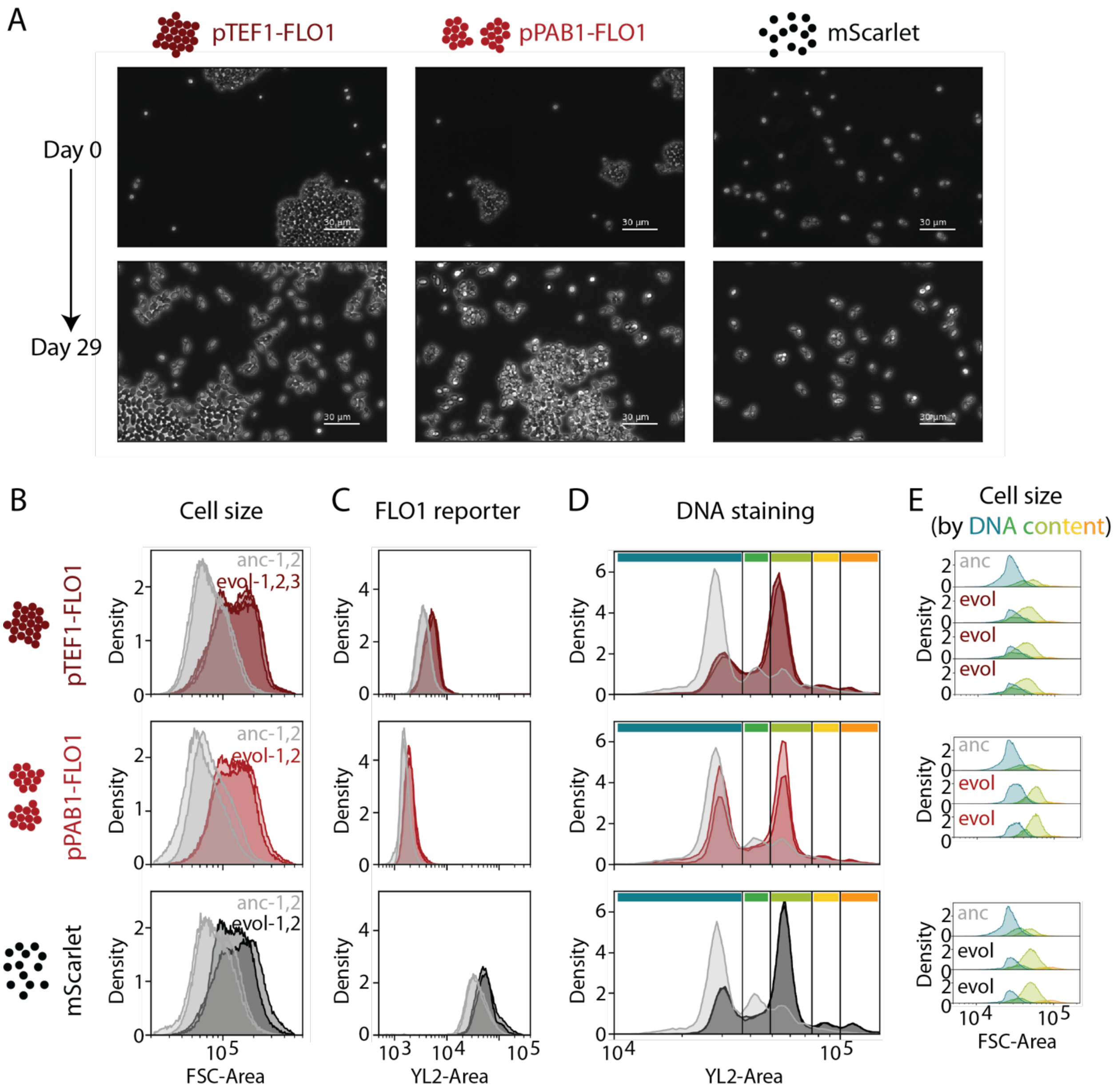


**Figure S2. Cell size increase in continuous culture occurs in all strains and is consistent with an increase in ploidy.**

(A) Over the course of continuous culture, both cell size and aspect ratio increased. This increase in aspect ratio also leads to less orderly packing in the aggregate, further increasing apparent aggregate size.

(B) Flow cytometry shows an increase in Forward Scatter (FSC), a proxy for cell size, for all continuously cultured strains, regardless of whether they flocculated. Replicates of the ancestral strain (e.g., anc-1, anc-2) are different overnight cultures of the same parental strain, while replicates of the evolved strains (e.g., evol-1, evol-2) are different eVOLVER vials, with an additional vial for the largest aggregates.

(C) mScarlet fluorescence also increased over the course of the experiment for both aggregating and non-aggregating strains, consistent with a possible increase in ploidy that is independent of FLO1 expression.
(D) Propidium iodide staining of ancestral and evolved strains shows an increase in the proportion of cells with higher DNA content. Overall, staining revealed 5 possible peaks of PI staining.
(E) Gating on the five peaks of PI staining shows that FSC increases with DNA content. These larger cells with higher DNA content are much more prevalent in evolved strains.

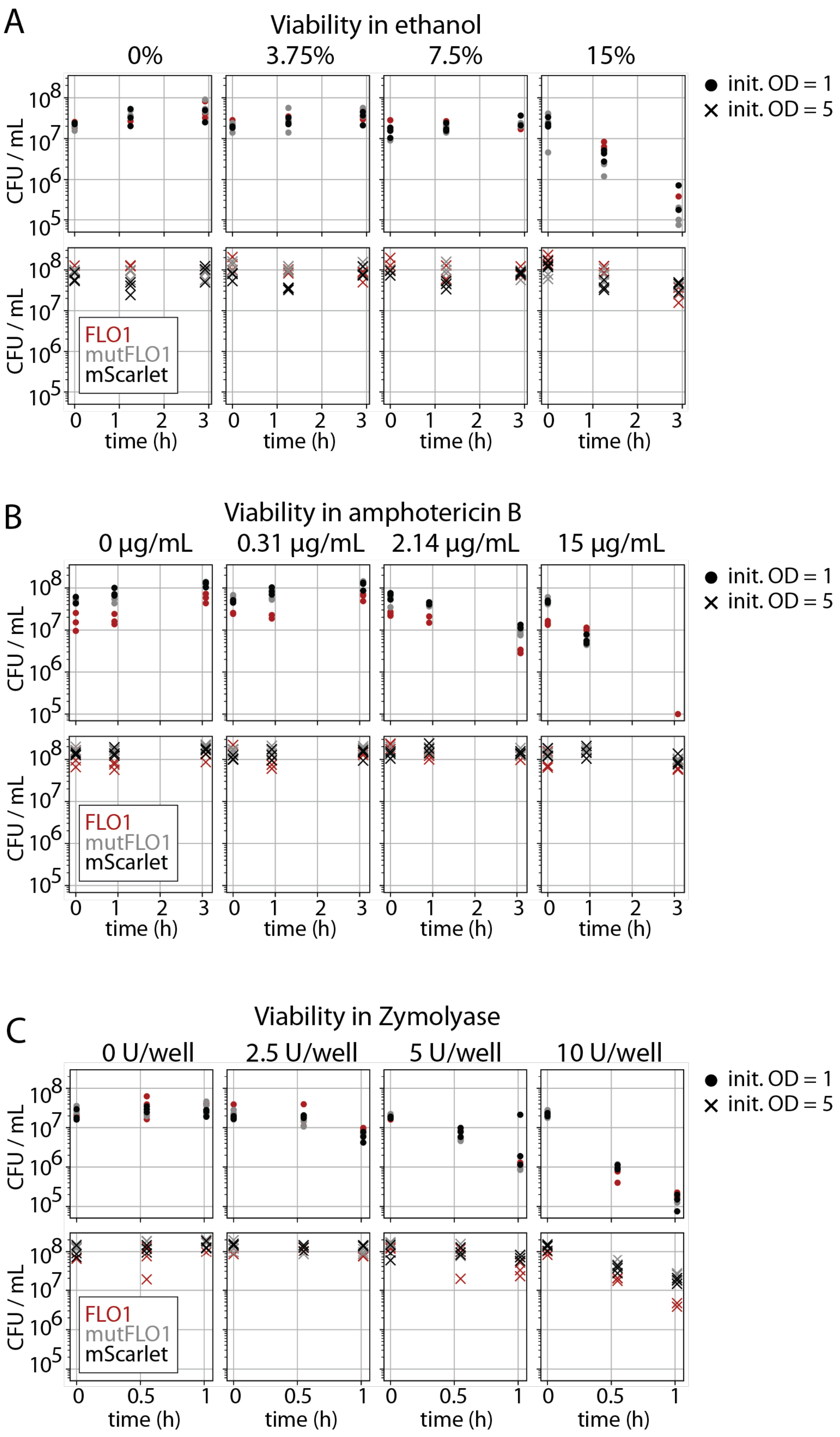
A
Viability in ethanol
0%
3.75%
7.5%
15%
init. OD = 1
init. OD = 5
CFU / mL
FLO1
mutFLO1
mScarlet
time (h)
B
Viability in amphotericin B
0 µg/mL
0.31 µg/mL
2.14 µg/mL
15 µg/mL
C
Viability in Zymolyase
0 U/well
2.5 U/well
5 U/well
10 U/well

**Figure S3. Viability over time in diverse chemical stresses is consistent with endpoint effects.**

Cell counts over time were estimated by fitting dilutions of countable colonies to a Poisson distribution. Cell counts at time 0 reflect the different initial densities in each culture. Dynamics are shown for ethanol (A), amphotericin B (B), and Zymolyase (C).

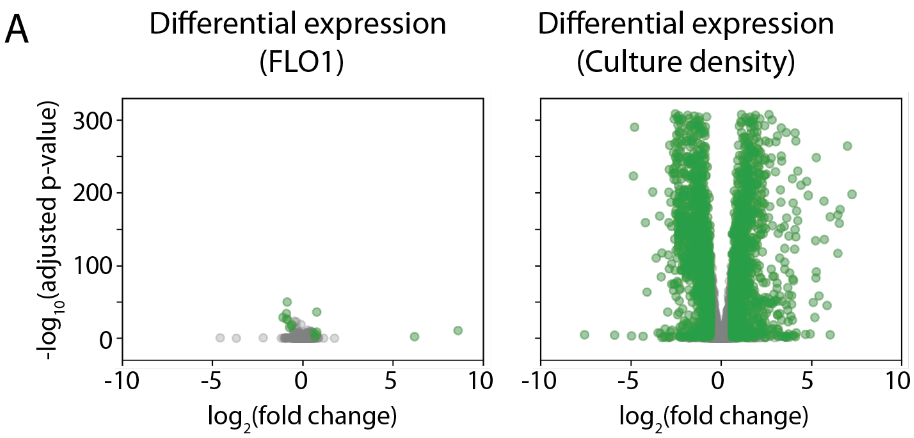


B

GO Term Analysis

| | GO Term | GO description | Relevant genes |
|---|---|---|---|
| 0 | GO:0015891 | siderophore transport | [SIT1, ARN1, ARN2, ARN2, FIT2, FIT3, FIT2] |
| 1 | GO:0006826 | iron ion transport | [ARN2, ARN1, SIT1, FIT2, FIT3, FRE5, FRE5] |
| 2 | GO:0006879 | intracellular iron ion homeostasis | [TIS11, SIT1, SIT1, ARN2, ARN1, FRE5, SIT1, ARN2] |
| 3 | GO:0033214 | siderophore-dependent iron import into cell | [ARN2, ARN1, SIT1, SIT1] |
| 4 | GO:0044718 | siderophore transmembrane transport | [ARN2, ARN1, SIT1, SIT1] |
| 5 | GO:0015343 | siderophore-iron transmembrane transporter act... | [ARN1, ARN2, ARN2, ARN2, ARN2, SIT1, ARN1] |

**Figure S4. Genes differentially regulated by FLO1 are enriched in GO terms for iron metabolism, though effect sizes are small.**

(A) $Log_2$ fold change of expression versus adjusted p-values show limited effect sizes of FLO1 expression, regardless of statistical significance, while culture density produces both large and statistically significant effects. Green dots were identified as significantly differentially expressed, with a significance threshold of 0.05 for the adjusted p-value and a minimum fold change of 1.5.

(B) Of the 16 genes differentially-expressed in synthetic flocs, six were enriched in GO terms related to iron transport. Notably, all six of these were downregulated when FLO1 was expressed.

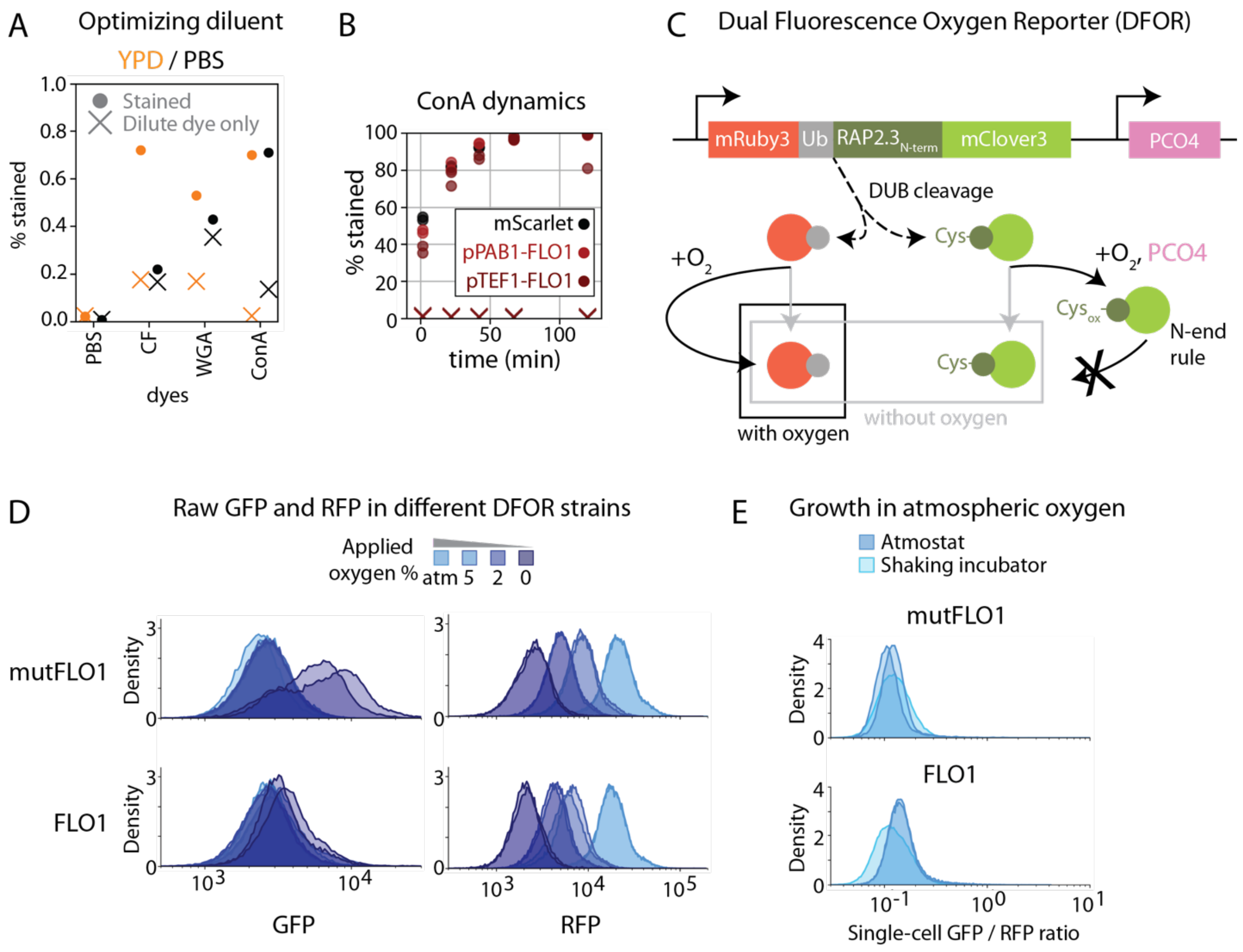


**Figure S5. Calibration of dye and oxygen diffusivity assays.**

(A) The mScarlet unicellular control was either stained and diluted tenfold (“Stained”) or only mixed with the tenfold diluted dye (“Dilute dye only”), using either YPD or PBS as the diluent. Relative to diluting in PBS, diluting in YPD reduced background while also preserving the degree of staining.

(B) Triplicate cultures were stained with ConA for different duration pulses before flow. While the dye was excluded from aggregates at early timepointst, all cultures asymptote to 100% staining in approximately one hour.

(C) The Dual Fluorescence Oxygen Reporter (DFOR) includes two fluorescent proteins fused by a ubiquitin monomer, which ensures the cleavage of the two proteins. Cleaved GFP contains an N-terminal plant domain (RAP2.3) whose initial cysteine can be oxidized. The second component is PCO4, a plant enzyme capable of this oxidation. This oxidation then targets the GFP for degradation via the N-end rule. In this way, RFP expression is constitutive, but GFP expression is oxygen-dependent.

(D) The DFOR construct displays different behavior in different background strains. In the strain expressing mutFLO1, GFP increases much more when oxygen is removed, and RFP values have a greater dynamic range. This underscores the greater reliability of computing the ratio of the two fluorophores and making comparisons within the same strain.

(E) The distribution of DFOR values for cells grown aerobically in the shaking incubator and in the eVOLVER with atmospheric oxygen are very similar, showing aeration in the eVOLVER atmostat module is likely similar to benchtop culture.

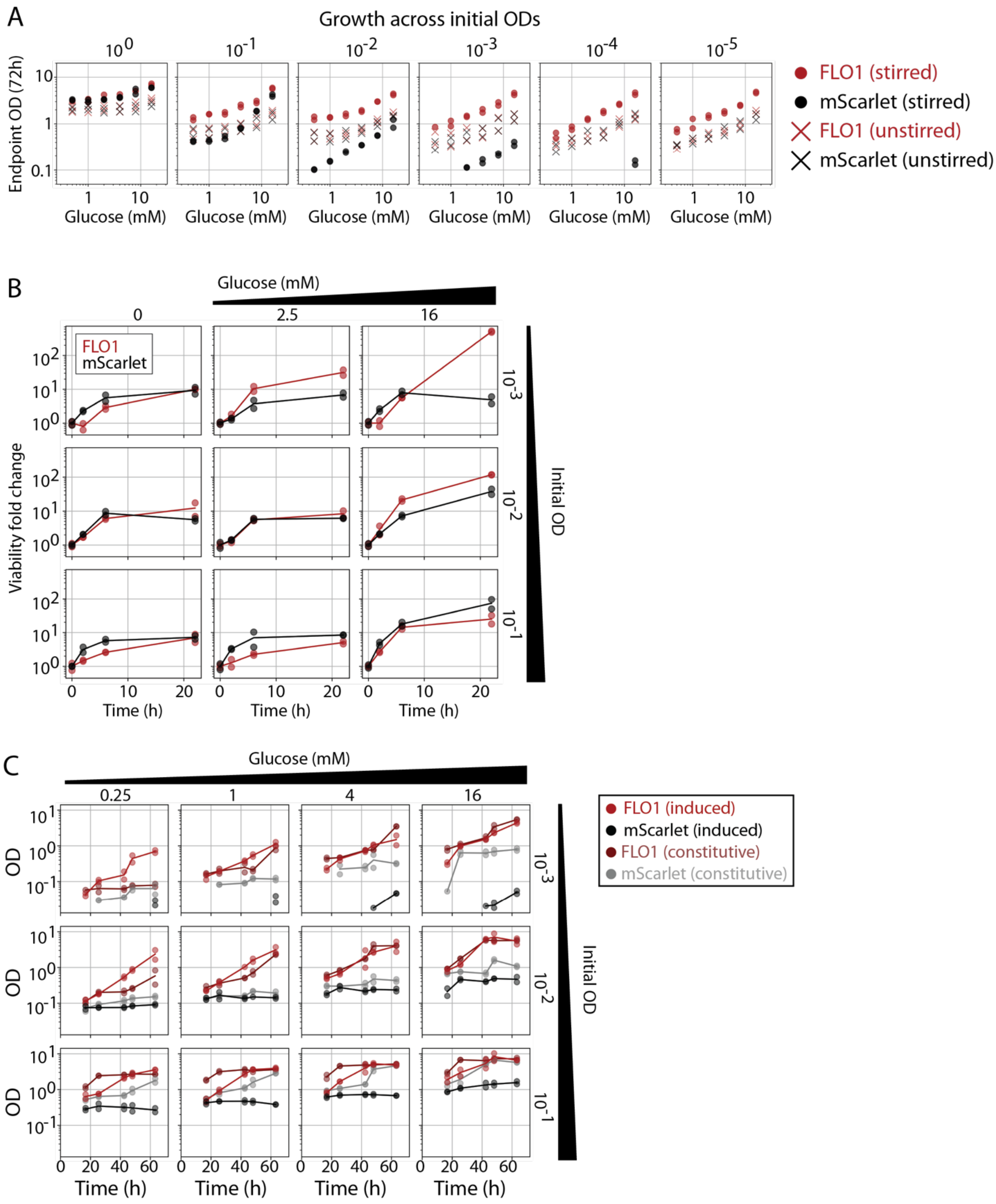


**Figure S6. Improved growth of aggregating strains is specific to stirred media and is also observed in strains constitutively expressing FLO1.**

(A) In limiting glucose, synthetic flocs grew better than the mScarlet strain only when cultures were stirred and the initial cell density was less than one. When unstirred, strains grow to the same extent, but stirring increases the growth of the FLO1 strain while decreasing growth of the mScarlet strain.

(B) Plating countable colonies reveals changes in viability at early timepoints, when OD was below the limit of detection of the plate reader. In the absence of glucose, strains do not lose viability but continue to grow, suggesting differences in endpoint OD reflect differences in growth rate, not differences in survival. The first ~10h of growth are similar in all strains, likely continued growth on intracellular stores from pregrowth in 2% glucose. Nonetheless, growth rate differences emerge at later timepoints.

(C) Synthetic flocs grew better than mScarlet strains, whether gene expressions as constitutive or inducible. However, the constitutive mScarlet strain had better growth than its inducible counterpart.

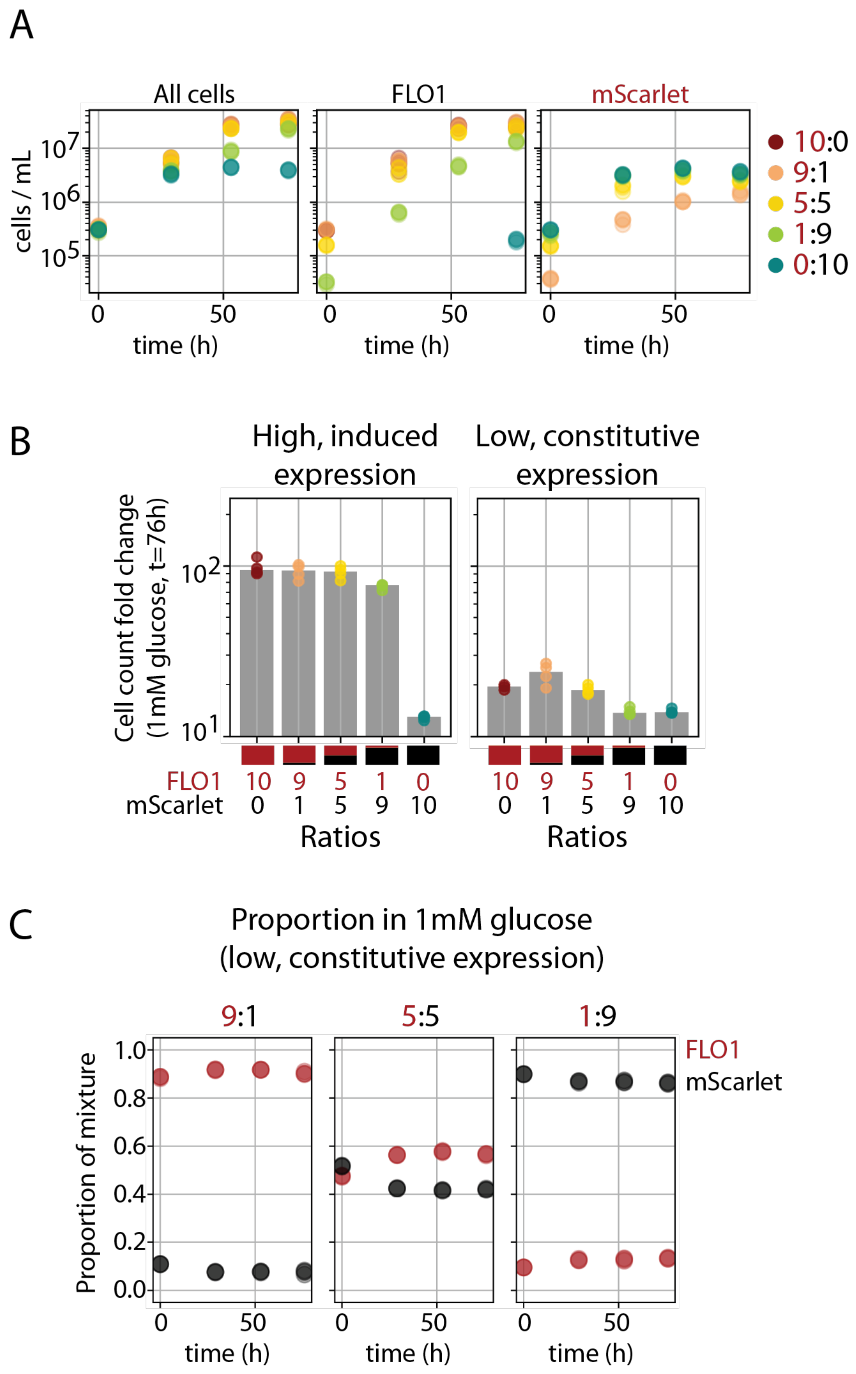


**Figure S7. Benefits of aggregation depend on FLO1 expression levels.**

(A) Tracking the proportion of synthetic flocs and the mScarlet strain during growth in 1mM glucose shows that mixtures diverge only after 24h. Synthetic flocs continue to grow and saturate at higher cell concentrations, while mScarletI-expressing cells in the same mixture grow slower and saturate sooner.

(B) Synthetic flocs and the mScarlet strain were mixed using either the inducible (as in Figure 7) or constitutive versions (as in Figure S6C). The benefit to synthetic flocs alone is smaller in the constitutive case (consistent with Figure S6C), and even smaller in mixtures. This is likely because expression is also lower in the constitutive strains (cf. Figure S1C), resulting in smaller aggregate sizes and therefore fewer benefits of nutrient sharing.

(C) Synthetic flocs do not outcompete the mScarlet strain as quickly in the constitutive case. However, this slower competition is consistent with the smaller benefit overall observed for constitutive strains.

### SUPPLEMENTAL TABLES

**Supplemental Table S1.** Plasmids used in this study.

**Supplemental Table S2.** Yeast strains used in this study.

### SUPPLEMENTAL NOTE 1

#### Estimating 3D aggregate size

Flattening aggregates into two dimensions preserves their size and permits more precise quantification. However, their 3D size is relevant for estimating diffusion limitations in liquid culture. To estimate the lower bound of 3D size, we assume that the volume of the cells (which we can estimate from the 2D image) forms 75% of the volume of the resulting aggregate, with cells packed at maximum density. (Random packing of cells into aggregates, or non-void volume ratio of 0.6, leads to similar estimates of aggregate sizes.)

$$V_{cells} = 0.75\, V_{agg}$$

To compute the volume of the aggregate, we assume it is a sphere.

$$V_{agg} = \frac{4}{3}\pi r_{agg}^3$$

To compute the volume of the cells, we convert the 2D area observed in microscope images to number of cells assuming the median radius of a yeast cell is 3μm. This corresponds to the reference line drawn on distributions of aggregate size and aligns well with median size observed for single cell controls. We then convert this number of cells into a spherical volume by multiplying by the volume of each cell.

$$V_{cells} = N_{cells}(\frac{4}{3}\pi r_{cell}^3) = \frac{A_{agg}}{\pi r_{cell}^2}\left(\frac{4}{3}\pi r_{cell}^3\right)$$

We then solve for the radius of the aggregate as a function of the observed 2D aggregate area.

$$r_{agg} = \sqrt[3]{\frac{A_{agg} r_{cell}}{0.75\pi}}$$

For some experiments, we do not have the exact aggregate area at the start of the experiment, but only the OD. We therefore use the aggregate area measured for the strain at the same OD in a different experiment. Figures 1 and S1 have paired OD and aggregate area measurements for the largest, constitutive strain (i.e., pTEF1-FLO1) at three different ODs. At OD = 2, the median area is $10^5$μm$^2$, corresponding to approximately 3500 cells and an aggregate radius of 50μm. At OD = 6, the median volume is $10^6$μm$^2$, corresponding to approximately 35000 cells and an aggregate radius of 100μm. Note that aggregate volume does not scale linearly with cell concentration (i.e., doubling the cell volume can more than double aggregate size), but this is not unexpected, as a null, chemical reaction kinetics model would assume cell-cell adhesion goes with the square of cell concentration. This 100μm is likely a lower bound still, as $10^6$μm$^2$ was the largest area we imaged for most aggregates (i.e., flattened aggregates could also be much larger).

We also estimated that large 2mm diameter aggregates reported in the literature could only arise in saturated cultures (Smukalla *et al*., 2008). Here, we go from the reported aggregate size to the number of cells and, to find a lower bound of cell culture density, assume the largest possible void volume (i.e. random packing).

$$V_{cells} = 0.6\, V_{agg}$$

Then we solve for $N_{cells}$.

$$N_{cells} = 0.6 \left(\frac{r_{agg}}{r_{cell}}\right)^3$$

For a 2mm radius aggregate made of 3μm radius cells, this is ~18x$10^7$ cells. Assuming these aggregates were composed of cells from 3mL cultures, this is ~6x$10^7$ cells / mL, or an OD of 6. We note that for our strains, such high density was sufficient to produce the kind of chemical protection and effects on gene expression that were previously attributed to flocculation.
